# Machine learning guided signal enrichment for ultrasensitive plasma tumor burden monitoring

**DOI:** 10.1101/2022.01.17.476508

**Authors:** Adam J. Widman, Minita Shah, Nadia Øgaard, Cole C. Khamnei, Amanda Frydendahl, Aditya Deshpande, Anushri Arora, Mingxuan Zhang, Daniel Halmos, Jake Bass, Theophile Langanay, Srinivas Rajagopalan, Zoe Steinsnyder, Will Liao, Mads Heilskov Rasmussen, Sarah Østrup Jensen, Jesper Nors, Christina Therkildsen, Jesus Sotelo, Ryan Brand, Ronak H. Shah, Alexandre Pellan Cheng, Colleen Maher, Lavinia Spain, Kate Krause, Dennie T. Frederick, Murtaza S. Malbari, Melissa Marton, Dina Manaa, Lara Winterkorn, Margaret K. Callahan, Genevieve Boland, Jedd D. Wolchok, Ashish Saxena, Samra Turajlic, Marcin Imielinski, Michael F. Berger, Nasser K. Altorki, Michael A. Postow, Nicolas Robine, Claus Lindbjerg Andersen, Dan A. Landau

## Abstract

In solid tumor oncology, circulating tumor DNA (ctDNA) is poised to transform care through accurate assessment of minimal residual disease (MRD) and therapeutic response monitoring. To overcome the sparsity of ctDNA fragments in low tumor fraction (TF) settings and increase MRD sensitivity, we previously leveraged genome-wide mutational integration through plasma whole genome sequencing (WGS). We now introduce MRD-EDGE, a composite machine learning-guided WGS ctDNA single nucleotide variant (SNV) and copy number variant (CNV) detection platform designed to increase signal enrichment. MRD-EDGE uses deep learning and a ctDNA-specific feature space to increase SNV signal to noise enrichment in WGS by 300X compared to our previous noise suppression platform MRDetect. MRD-EDGE also reduces the degree of aneuploidy needed for ultrasensitive CNV detection through WGS from 1Gb to 200Mb, thereby expanding its applicability to a wider range of solid tumors. We harness the improved performance to track changes in tumor burden in response to neoadjuvant immunotherapy in non-small cell lung cancer and demonstrate ctDNA shedding in precancerous colorectal adenomas. Finally, the radical signal to noise enrichment in MRD-EDGE enables *de novo* mutation calling in melanoma without matched tumor, yielding clinically informative TF monitoring for patients on immune checkpoint inhibition.

## INTRODUCTION

Liquid biopsy offers to reshape cancer care through the noninvasive detection and monitoring of plasma circulating tumor DNA (ctDNA). The clinical potential of this emerging biomarker has fostered a diversity of approaches designed to capture ctDNA signal from the broader plasma cell-free DNA (cfDNA) pool, including mutation-based approaches such as deep targeted panels^1–8^, approaches centered around cfDNA fragmentation patterns and coverage footprints^9–11^, and strategies focused on cancer-specific methylation and epigenetic patterns^12–16^. Clinically, ctDNA mutational profiling is increasingly used in high tumor fraction (TF) disease (e.g., non-invasive mutation detection to guide targeted therapy^1,17,18^).

Extensive recent efforts have focused on extending the use of ctDNA mutation detection to low TF settings such as therapeutic response monitoring or the assessment of minimal residual disease (MRD). The detection of residual ctDNA after surgical or non-surgical interventions could enable precision tailoring of treatment, offering treatment intensification or de-escalation based on MRD status. To overcome the inherent ctDNA signal sparsity in low TF settings such as MRD, many have employed deep targeted sequencing to capture mutations from tumor-informed bespoke panels^19,20^ or common cancer driver genes^4,5,8,21^. Missed detections, however, remain prevalent in current assays. For example, MRD identified via bespoke panels in urothelial carcinoma is strongly prognostic of disease recurrence, though up to 40% of ctDNA-negative patients experienced relapse^19^. Similar ‘false negatives’ were seen in breast^5^ and colorectal cancer^22–24^, suggesting that further improvement in sensitivity is needed.

We and others^25–28^ have previously demonstrated that sensitivity barriers in deep targeted panels arise from the limited number of ctDNA fragments recovered at targeted loci. Even with ideal error suppression and ultra-deep sequencing, a somatic mutation cannot be observed if it is not sampled in the limited plasma volume collected in routine testing, which imposes a hard barrier on effective coverage depth. Sensitivity is therefore tied to the limited number of genome equivalents (GE) in a plasma sample (typically 1,000s per mL^29^), and when TF is below harvested GEs, MRD detection is diminished. Targeted approaches have sought to overcome this limitation by increasing the number of panel-covered mutations to dozens^3,8,19–21^ or even 100s^25^ or enriching for biological features of ctDNA such as altered fragment size^7,30^.

We previously proposed an alternative approach in which breadth of sequencing could supplant depth of sequencing via integration of thousands of single nucleotide variants (SNVs) and copy number variants (CNVs) across the cancer genome^28^. We implemented whole genome sequencing (WGS) of plasma and matched tumor for enhanced MRD signal recovery in colorectal cancer (CRC) and lung adenocarcinoma (LUAD). Our accompanying denoising approach MRDetect enabled the detection of plasma TFs as low as 1*10^-5^ and identified postoperative MRD linked to early disease recurrence^28^, supporting WGS as a viable strategy for MRD detection.

WGS allows for increased signal recovery at the expense of increased sequencing noise, yet denoising tools such as high sequencing depth and molecular tags leveraged by deep targeted panels are not typically deployed in the WGS setting. In our previous MRDetect work, we designed a support vector machine approach to identify patterns specific to WGS sequencing error and suppress low quality SNV artifacts. Herein we posit that learning patterns specific to ctDNA mutagenesis can offer signal enrichment to complement suppression of sequencing error. We developed MRD-EDGE (**E**nhanced ct**D**NA **G**enomewide signal **E**nrichment), which integrates complementary signal from SNVs and CNVs to increase ctDNA signal enrichment in plasma WGS. For SNVs, MRD-EDGE uses deep learning to integrate the myriad local and regional properties of somatic mutations to identify ctDNA mutations among sequencing error. For CNVs, MRD-EDGE uses machine learning-based denoising and an expanded feature space including fragmentomics and allelic frequency of germline single nucleotide polymorphisms (SNPs) to enable ultrasensitive ctDNA detection at lower degrees of aneuploidy than MRDetect. The increased performance of MRD-EDGE enabled ultrasensitive MRD and tumor burden monitoring in tumor-informed settings, as well as the detection of ctDNA shedding from precancerous colorectal adenomas. Further, the signal to noise enrichment from MRD-EDGE enabled *de novo* (nontumor-informed) detection of melanoma ctDNA SNVs at sensitivity on par with tumor-informed targeted panels. We demonstrate the clinical utility of this *de novo* approach by using plasma ctDNA response to immune checkpoint inhibition (ICI) to predict long-term treatment outcomes.

## RESULTS

### Deep learning integrates mutagenesis features to distinguish ctDNA SNVs from sequencing error

A prominent obstacle to WGS-based detection of ctDNA SNVs is distinguishing true tumor mutations from far more abundant sequencing error. In our previous work^28^, we developed an error suppression framework that operates at the individual fragment (rather than locus) level. This significant departure from traditional consensus mutation callers was driven by the expectation that in standard WGS coverage (e.g., 30X) of low TF samples (e.g., TF < 1:1000), at best only a single supporting fragment will be detected for any given mutation. A support vector machine (SVM) classification framework was applied to exclude error associated with lower quality sequencing metrics including variant base quality (VBQ), mean read base quality (MRBQ), variant position in read (PIR), and paired-read mutation overlap. Focused solely on eliminating sequencing error, the classifier was trained on reads with germline SNPs (true labels) vs. reads with sequencing errors (false labels).

We posited that signal to noise enrichment may emerge not only from characterizing features specific to sequencing errors (decreasing noise), but also from learning features indicative of true ctDNA mutations (increasing signal). Learning features specific to ctDNA required a rethinking of our machine learning training paradigm, as germline SNPs can no longer serve as a source for true (positive) labels. Instead, we leveraged cfDNA samples with high TF (range 9-24%, **Supplementary Table 2**) across three common cancer types with high mutational burden: melanoma, LUAD, and colorectal cancer. These high TF plasma samples (range *n*=2-4) provided an abundant (51,160 to 270,648, **Supplementary Table 2**) source of fragments enriched with somatic mutations (true labels) from which to develop a ctDNA SNV feature space. Our ctDNA SNVs were compared to cfDNA fragments containing sequencing errors drawn from controls (range *n*=4-5) without a known malignancy (**Supplementary Table 2** and Methods). To ensure that classification is optimized to detect more subtle differences between signal and noise, we implemented a set of quality filters to remove germline SNPs, recurrent plasma WGS artifacts, and variants with low base or mapping quality scores (**Supplementary Table 3** and Methods).

After obtaining a large, pre-filtered training corpus of ctDNA SNVs and cfDNA SNV artifacts, we next explored a broader feature space to help distinguish the two. First, single base substitutions (SBS) sequence patterns are closely associated with cancers driven by distinct mutational processes^31,32^ such as SBS4 signature (tobacco exposure) in LUAD or SBS6 (ultraviolet light) in melanoma. Second, ctDNA has been associated with shorter fragment size^30,33,34^. Third, SNVs are overrepresented in distinct locations within the genome, including a predilection for quiescent chromatin and late replicating regions^35–38^, allowing for inference of the local (e.g., 20Kb) mutation likelihood. This evaluation allowed us to identify informative features with varying contribution across tumor types (**Fig 1b**, **Extended Data 1a**, **Supplementary Table 3**).

**Figure 1:**
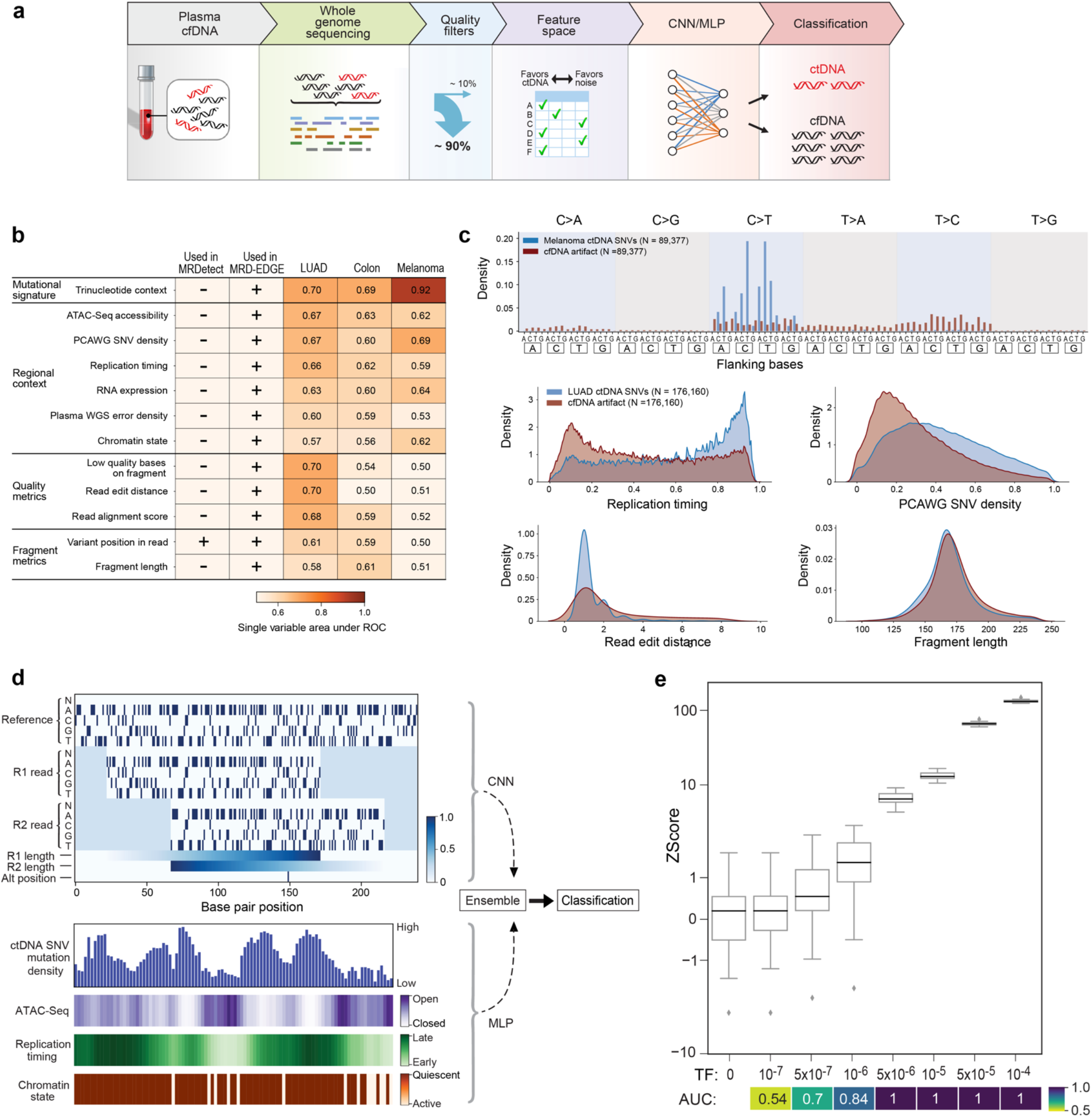
Application of disease-specific deep learning classifier to distinguish ctDNA SNV fragments from cfDNA artifacts. **a)** Illustration of whole genome sequencing (WGS)-based ctDNA single nucleotide variant (SNV) detection in plasma with MRD-EDGE. Healthy cfDNA and ctDNA are admixed in the plasma pool. Both cfDNA and ctDNA are subjected to WGS, and SNVs are identified against the reference genome and subjected to quality pre-filters designed to reduce artifact from sequencing error and germline variants. A complex feature space designed to distinguish ctDNA signal from cfDNA noise serves as input to a deep learning neural network, where fragments containing SNVs are classified as ctDNA or cfDNA with sequencing artifacts. **b)** Heatmap of selected *post-filter* model features and the single variable area under the receiver operating curve (svAUC) between individual features and label (ctDNA or cfDNA) in LUAD, CRC, and melanoma. In this comparison, ctDNA SNV fragments and cfDNA SNV artifacts are drawn from within the same plasma sample to remove potential inter-sample biases when establishing predictive capacity of individual features. For categorical features, AUC was assessed on a held-out validation set of fragments after a linear classifier was trained to predict positive or negative label based on one-hot encoded categorical features. Features are annotated with whether they are used in MRDetect or MRD-EDGE. **c)** Selected feature density plots for post-filter ctDNA and cfDNA SNV artifacts: trinucleotide context, replication timing^37^, PCAWG^81^ tumor SNV mutation density, read edit distance, and fragment length. **d)** (top) Illustration of the fragment tensor, an 18×240 matrix encoding of the reference sequence, R1 and R2 read pairs (including padding where reads do not overlap the reference sequence), R1 read length and R2 read length, and the position of the SNV in the fragment (‘Alt position’). The fragment architecture allows for integration of fragment-specific features such as trinucleotide context, fragment length, and edit distance, among others. The fragment tensor is passed as input to a convolutional neural network. (bottom) Illustration of the relationship between regional features and local ctDNA SNV mutation density at the chromosome level. Disease-specific inaccessible^82^ and quiescent^83^ genomic regions, as well as late replicating regions^37^, are associated with somatic mutagenesis as represented by increased density of tumor-confirmed ctDNA SNVs. Regional features (**Supplementary Table 3**) are encoded as tabular values and passed as input to a multilayer perceptron. An ensemble classifier takes input from both the fragment and regional models to determine the likelihood that each fragment is ctDNA or cfDNA SNV artifact. **e)** *In silico* studies of cfDNA from the metastatic cutaneous melanoma sample MEL-01 mixed into cfDNA from a healthy plasma sample (‘C-16’) at mixing fractions TF = 10^-7^–10^-4^ at 16X depth, performed in 20 technical replicates with independent sampling seeds. Tumor-informed MRD-EDGE enables sensitive TF detection as measured by Z score against unmixed control plasma (TF=0, *n=*20 randomly chosen replicates) as low as TF=5×10^-7^ (AUC 0.70). Box plots represent median, lower and upper quartiles; whiskers correspond to 1.5 x IQR. An AUC heatmap benchmarks detection sensitivity vs. TF=0 at different mixed TFs. IQR, interquartile range.

To integrate this expanded feature set for optimal classification, we reasoned that neural networks would best serve the size of our training sets (100,000s of fragments) and the underlying feature complexity. We developed a two-dimensional representation of a cfDNA fragment (**Fig 1d**, top and Methods) to capture fragment-level features such as SBS, fragment length, and quality metrics like read edit distance and PIR. In parallel, a second model architecture was designed to capture regional context, whereby each SNV-containing fragment is scored based on salient regional features associated with mutation frequency (**Fig 1d**, bottom). For example, a fragment can be annotated with the local density of melanoma tumor SNVs in a 20Kb interval surrounding the candidate SNV (Methods, **Supplementary Table 3** for a full list of features by cancer type). We combined our fragment and regional architectures as inputs to an ensemble model featuring a convolutional neural network (fragment CNN) for our fragment architecture and a multilayer perceptron (regional MLP) for our regional architecture. This ensemble model uses a sigmoid activation function to output a score between 0 and 1 to indicate the likelihood that a candidate SNV is either cfDNA sequencing error or a ctDNA mutation. Our ensemble model outperformed both the fragment and region models individually and other machine learning architectures in a melanoma validation plasma sample (‘MEL-01’) held out from training and paired with SNV artifacts from healthy control plasma (**Extended Data 1b**, Supplementary Table 2). We note that our deep learning methods were applied to a more stringent classification task than in our previous work, as we applied our classifier to heavily pre-filtered fragments in which the majority of low quality cfDNA sequencing errors were excluded (mean 92.8%, range 91.2%-93.6%). In this context, we found that our classification method yielded area under the receiver operating curves (AUCs) at the fragment level of 0.95 (95%: 0.94-0.95) in melanoma, 0.87 (0.86-0.88) in LUAD, and 0.84 (0.83-0.84) in colorectal cancer in validation plasma samples held out from training (**Extended Data 1c**, Supplementary Table 2).

We next sought to benchmark our platform’s enrichment capacity in the tumor-informed setting, in which a patient-specific mutational compendia drawn from resected tumor tissue is used to nominate SNVs for classification. We used tumor-confirmed ctDNA SNVs from MEL-01 admixed with SNV artifacts drawn from 6 healthy control plasma samples that were held out from model training (‘Melanoma held-out validation fragments’, **Supplementary Table 2**). First, we measured signal to noise enrichment for the pipeline as a whole and at individual stages (**Extended Data 1d**). Given the higher likelihood of a true positive in the tumor-informed setting, we used a balanced classification threshold (0.5) on the final ensemble model to classify ctDNA signal from noise. In a matched analysis in which both platforms were applied to the same data, we found a higher signal to noise (S2N) enrichment for MRD-EDGE (mean 118-fold, range 100-153 fold) compared to MRDetect (mean 8.3-fold, range 8-9 fold), which translates to a mean additional 14-fold S2N enrichment (range 12-18 fold).

We next evaluated the lower limit of detection (LLOD) for our tumor-informed MRD-EDGE classifier in *in silico* TF admixtures (TFs 10^-4^-10^-7^, *n*=20 *in silico* admixture replicates, Methods) using reads from MEL-01 mixed into control cfDNA from an individual (‘C-16’) with no known cancer (**Fig 1e**). When compared to the noise distribution in randomly chosen TF=0 replicates, we found higher performance even in the parts per million range and below (AUC of 0.84 at TF 1*10^-6^ and 0.7 at 5*10^-7^ for MRD-EDGE, compared to 0.77 and 0.65 for MRDetect, respectively).

### Advanced denoising and an enriched feature space enable enhanced CNV-based ctDNA detection

Aneuploidy is observed in the vast majority of solid tumors and is a prominent hallmark of the cancer genome^39^. We have shown that MRDetect-based CNV detection can monitor disease burden in cancers with a high degree of aneuploidy but low SNV mutation burden^28^. MRDetect sought to identify plasma read depth skews corresponding to matched tumor-informed CNV profiles to measure MRD in CRC and LUAD. While our results demonstrated a 2 order of magnitude improvement in sensitivity compared to leading CNV-based ctDNA algorithms^10,28^, we required substantial aneuploidy (>1Gb altered genome) to detect TFs of 5*10^-5^.

We reasoned that detection of subtle read depth skews related to low TF ctDNA may be hindered by biases that arise from sample preparation (e.g., GC bias), alignment (e.g., variable mapping), and biological factors (e.g., replication timing). These biases can introduce distortions (‘waviness’) in read depth signal which interfere with CNV estimation in both tumors and plasma^40^. To correct for such biases, we developed a machine-learning guided CNV denoising platform for use in plasma WGS. Our plasma read depth classifier uses robust principal component analysis (rPCA) trained on a panel of normal samples (PON) to correct read depth distortions due to background artifacts related to assay, batch, and recurrent noise (Methods).

To evaluate the performance of ctDNA detection with our enhanced read depth classifier, we admixed *in silico* reads from a pretreatment high burden melanoma plasma sample with a high degree of aneuploidy (‘AD-12’, TF 17% with 1.6 GB of total aneuploidy, **Supplementary Table 2**) into a posttreatment sample from the same patient following a major response to immunotherapy, varying the TF admixtures (range 10^-3^–10^-6^; *n*=50 technical admixing replicates with random independent seeds). We identified signal from read depth skews at TF admixtures as low as 1*10^-5^ (**Fig 2b**). Directional skew signal from copy neutral regions in the matched tumor served as a negative control (**Extended Data 2d**).

**Figure 2:**
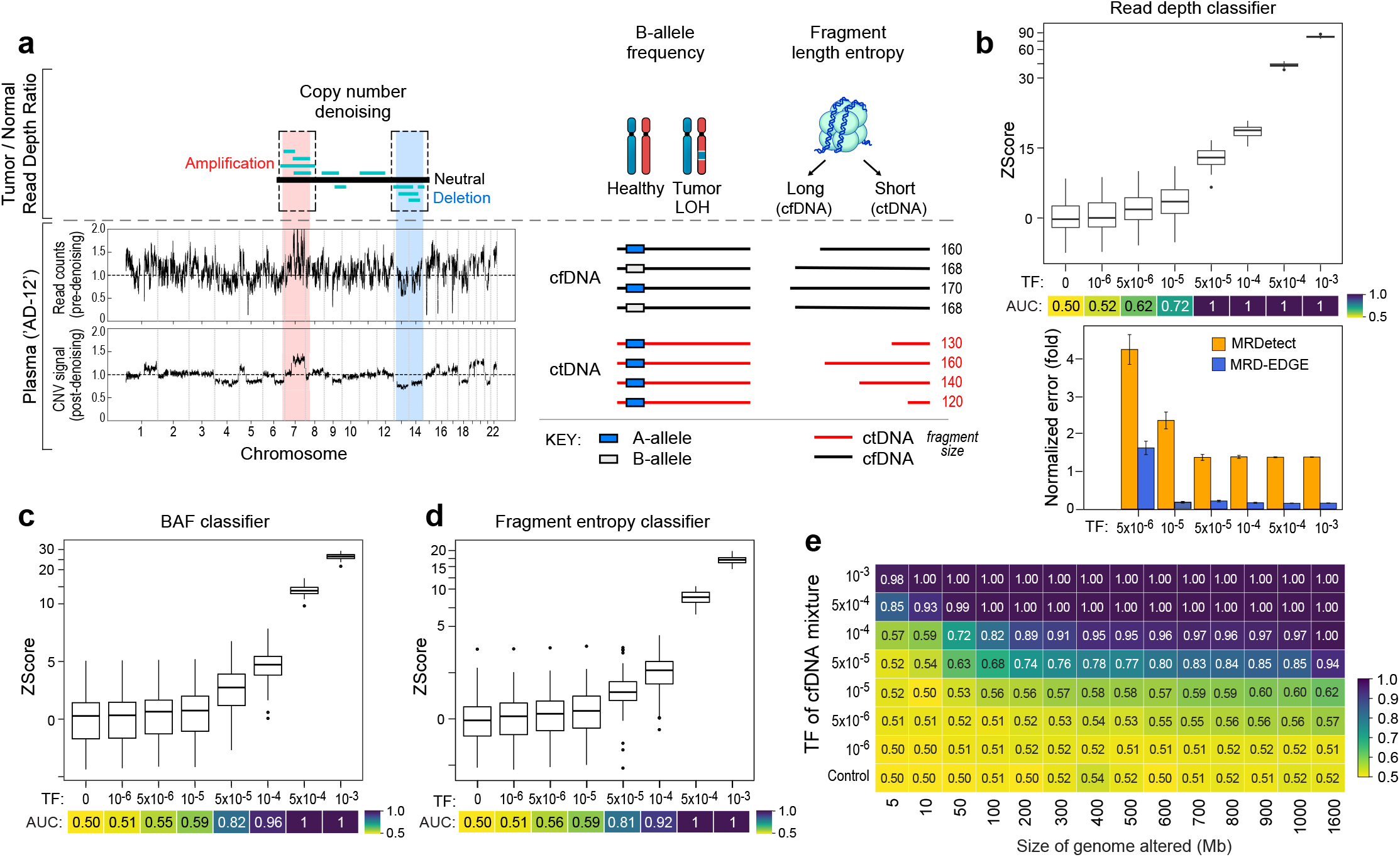
Machine learning-based error suppression and additional features enhance plasma WGS-based CNV detection sensitivity. **a)** (left) Illustration depicting use of copy number denoising for inference of plasma read depth. (top left) Patientspecific CNV segments are selected through the comparison of tumor and germline WGS. In plasma, these CNV segments may be obscured within noisy raw read depth profiles (middle left). Machine-learning guided denoising through use of a panel of normal samples (PON) drawn from healthy control plasma samples removes recurrent background noise to produce denoised plasma read depth profiles (bottom left). Plasma samples used in the PON are subsequently excluded from downstream CNV analysis. (middle) Loss of heterozygosity (LOH) results in replacement of heterozygous single nucleotide polymorphisms (SNPs) with homozygous variants and can be measured via changes in the B-allele frequency of SNPs in cfDNA. (right) Increased or decreased fragment length heterogeneity is expected in regions of tumor amplifications or deletions, respectively, due to varying contribution of ctDNA (shorter fragment size) to the plasma cfDNA pool. Fragment length heterogeneity is measured through Shannon’s entropy of fragment insert sizes. Fragment entropy signal is aggregated based on matched tumor amplifications (positive signal) or deletions (negative signal). **b-e)** *In silico* mixing studies of admixed high and low TF samples from the melanoma patient AD-12. Pretreatment plasma (TF = 17%) was mixed into posttreatment plasma (TF undetectable following a major response to immunotherapy) in 50 replicates. Admixtures model tumor fractions of 10^-6^–10^-3^. Box plots represent median, lower and upper quartiles; whiskers correspond to 1.5 x IQR. An AUC heatmap demonstrates detection performance at the different admixed TFs vs. negative controls (TF=0, *n*=25 replicates used to generate the noise distribution and *n*=25 used to benchmark performance) as measured by Z score. **b)** (top) Copy number denoising with the read depth classifier demonstrates detection sensitivity above TF=0 as low as 1*10^-5^ (AUC 0.72). (bottom) Normalized error at different mixed TFs between MRD-EDGE read depth classifier and MRDetect. Error is measured as 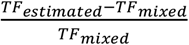. **c-d)** SNP BAF **(c)** and fragment length entropy **(d)** classifiers demonstrate Z score detection sensitivity at 5*10^-5^ (AUC 0.82 and 0.81, respectively). **e)** Empiric measurement of the MRD-EDGE lower limit of detection for the combined feature set as a function of the CNV load and admixture modeled TF. Sensitive detection (AUC 0.74) is observed at TF = 5*10^-5^ at 200 Mb. IQR, interquartile range. AUC, area under the receiver operating curve.

In addition to enhanced denoising of read depth skews, we reasoned that loss of heterozygosity (LOH) can serve as an important additional source of CNV signal. Copy neutral LOH cannot be captured by read depth skews but can be nonetheless measured through allelic imbalances in germline SNPs in plasma. Here, inference of the major allele in genomic regions affected by LOH is derived from tumor WGS^41–42^, and perturbations of the B-allele frequency (BAF) in plasma are indicative of ctDNA contribution to the plasma cfDNA pool (**Fig 2a**). To leverage LOH signal, plasma SNPs are aggregated in large genomic windows (1Mbp) and assessed for window-wide allelic imbalance. To account for underlying biases and mosaicism within the cfDNA pool, BAF values are compared both to the expected contribution of 0.5 and to the underlying peripheral blood mononuclear cell (PBMC) BAF reference^43^ (Methods), and quality filters are used to exclude aberrant signal due to low coverage and bias from PBMC (**Extended Data 2f**). Benchmarking of our BAF classifier in the same *in silico* admixtures yielded allelic imbalance signal in LOH regions in TF admixtures as low as 5*10^-5^ (**Fig 2c**).

Finally, we sought to leverage well-characterized abnormal ctDNA fragmentation patterns^9,33,34,44,45^ as an additional source of aneuploidy signal. ctDNA is associated with shorter and more heterogenous fragment lengths than normal cfDNA^9,44^. We therefore measured fragment length entropy (measured as Shannon’s entropy), a marker of heterogenous fragment lengths in cfDNA, in plasma WGS segments matched to amplifications and deletions in tumor. While existing approaches have sought to recognize altered fragmentation profiles inherently or compared to control (non-cancer) plasma^9,46^, in our fragment entropy classifier, use of matched tumor tissue enables the cfDNA fragment pool in neutral plasma regions to act as an internal control. Fragment lengths in matched CNV segments can be assessed in comparison to copy-neutral segments rather than to an absolute baseline, removing confounding from baseline fragment length biases at the sample level. We then measure the entropy contributions from amplifications (greater plasma cfDNA entropy due to a larger contribution of ctDNA fragments) and deletions (less plasma cfDNA fragment entropy) to harness signal. In our *in silico* admixtures, our fragment entropy classifier identified signal in TFs as low as 5*10^-5^ (**Fig 2d**, Methods). To demonstrate sensitivity across cancer types, we also benchmarked our CNV features in TF admixtures derived from pre- and postoperative plasma from a patient with early-stage non-small cell lung cancer (NSCLC) and found similar performance (**Extended Data 2a-c**).

The three CNV classifiers - read depth, BAF, and fragment entropy - gather independent and complementary sources of CNV signal. MRD-EDGE combines signal from these classifiers as independent inputs at the sample level to comprehensively assess for plasma TF (Methods). Because the aneuploidy signal in plasma WGS is a function of both the proportion of the cancer genome affected by aneuploidy and the TF, we evaluated classifier performance by downsampling both the TF (as above in **Fig 2b-d**) and the cumulative size of CNV segments to characterize a LLOD matrix (**Fig 2e**). Classifier performance, as expected, improved with increased aneuploidy. While MRDetect required 1 Gb of aneuploidy^28^ for a LLOD of 5*10^-5^, MRD-EDGE achieved an LLOD of 5*10^-5^ (AUC 0.74) with only 200Mb of aneuploidy, which would extend applicability to many more solid tumors (**Extended Data 3**).

### MRD-EDGE yields high performance in tumor-informed detection of early-stage colorectal cancer and postoperative MRD

To evaluate MRD-EDGE in the tumor-informed early-stage cancer setting, we tested the platform on our previously reported^28^ clinical cohort of plasma samples from patients with CRC (*n*=19, including 6 with microsatellite instability), compared with exposure matched controls without known cancer (*n*=34, ‘Control Cohort A’) and from the same sequencing platform (Illumina HiSeq X). Here, SNVs and CNVs from resected tumors form a patient-specific mutational compendia, which is then used to assess for ctDNA in pre- and postoperative plasma and to form noise (sequencing error) distributions in healthy control plasma. Z scores of patient plasma signal are derived from control plasma noise distributions and used to assess for ctDNA detection in both the MRD-EDGE SNV and CNV platforms independently (Methods). The Z score detection threshold was set at 90% specificity against control plasma in the receiver operating curve (ROC) analysis, and a positive ctDNA detection was defined as patient plasma SNV or CNV Z score above this threshold.

In our early-stage CRC cohort, area under the curve (AUC) for preoperative ctDNA SNV detection with MRD-EDGE was 1.00 (95% CI: 0.99 to 1.00) and sensitivity was 100% at 90% specificity (compared with MRDetect AUC 0.97, 95% CI: 0.91 - 1.00, 95% sensitivity at 90% specificity, **Fig 3a**). A cross-patient analysis, where the patient-specific mutational compendia was compared between matched and unmatched plasma, showed similar performance (**Extended Data 4a**). We note that our MRD-EDGE CRC SNV classifier was trained on high burden plasma sequenced with a different sequencing platform and at a different facility than the one used for the early-stage CRC samples (Illumina NovaSeq v1.5, Aarhus University, Denmark vs. Illumina HiSeq X, New York Genome Center, **Supplementary Table 1**), demonstrating generalizability across platforms. MRD-EDGE for CNVs was applied independently to this preoperative cohort and demonstrated improved performance (AUC = 0.82, 95% CI 0.71 - 0.91, 61% sensitivity at 90% specificity) compared to MRDetect (AUC = 0.73, 95% CI: 0.59 - 0.83, sensitivity = 40% at 90% specificity, **Fig 3b**). Moreover, the ability to evaluate copy neutral LOH in MRD-EDGE allowed us to apply CNV-based detection to 18 / 19 samples in this CRC cohort compared to 15 / 19 samples with MRDetect.

**Figure 3:**
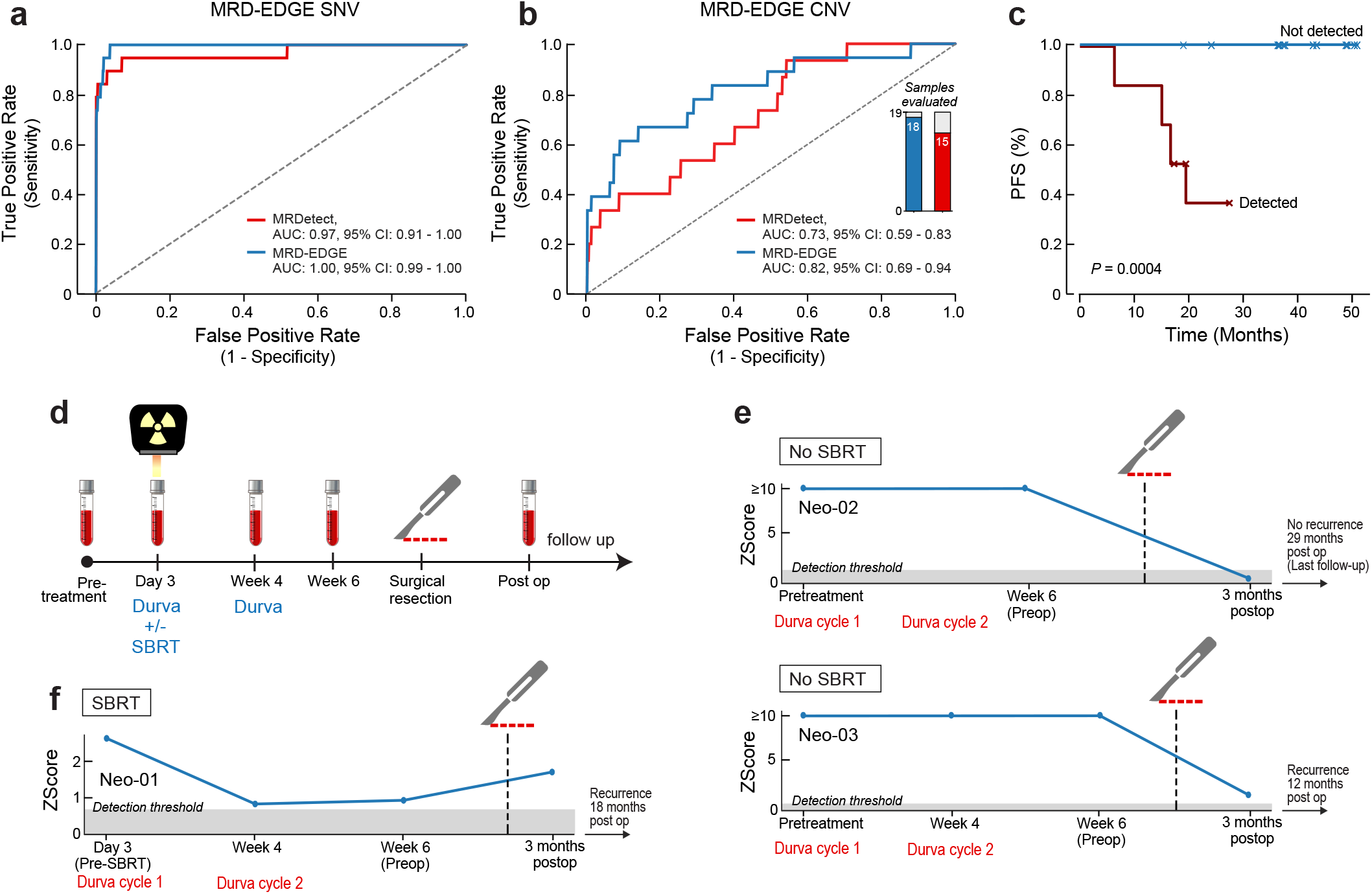
Detection of postoperative colorectal ctDNA and tracking neoadjuvant response to immune checkpoint inhibition and radiation in non-small cell lung cancer. **a)** ROC analysis on preoperative colorectal SNV mutational compendia for MRD-EDGE (blue) and MRDetect (red). Preoperative plasma samples (*n*=19) were used as the true label, and the panel of control plasma samples against all patient mutational compendia (*n*=646; 19 mutational compendia assessed across 34 control samples from Control Cohort A) was used as the false label. **b)** ROC analysis on preoperative colorectal CNV mutational compendia for MRD-EDGE (blue) and MRDetect (red) methods. Preoperative plasma samples (*n*=18, 1 sample excluded due to insufficient aneuploidy) were used as the true label, and the panel of control plasma samples against all patient mutational compendia (*n*=180; 18 mutational compendia assessed across 10 control samples from Control Cohort A) was used as the false label. Twenty-four samples from Control Cohort A were included in the read depth classifier panel of normal samples (PON, **Fig 2a**) and were held out from the CNV ROC analysis. **c)** Kaplan-Meier disease-free survival analysis was done over all patients with detected (*n*=9) and non-detected (*n*=10) postoperative ctDNA. Postoperative ctDNA detection shows association with shorter recurrence-free survival (two-sided log-rank test). **d)** Illustration of the neoadjuvant non-small cell lung cancer (NSCLC) clinical treatment protocol^50^. Plasma TF is tracked throughout the preoperative period to evaluate for response to SBRT and ICI therapy and after surgery to detect the presence of MRD. The detection threshold for MRD reflects 90% specificity in an independent cohort of preoperative patients with early-stage LUAD evaluated previousuly^28^ (**Extended Data 4c**). **e)** serial tumor burden monitoring on neoadjuvant immunotherapy with MRD-EDGE in 2 NSCLC patients on ICI therapy (no SBRT). Tumor burden estimates are measured as the Z score of the patient-specific mutational compendia against healthy control plasma. In both patients, unchanged plasma TF Z score demonstrates lack of response to ICI prior to surgery. (top) Upon surgical resection, there is no evidence of MRD and no recurrence at 29 months (patient Neo-02). (bottom) Upon surgical resection, plasma TF is above the detection threshold indicative of MRD, and disease recurrence is seen at 12 months postoperatively (patient Neo-03). **f**) demonstration of plasma TF decrease following radiation in a patient who was randomized to receive SBRT. ctDNA remains detectable following SBRT, and tumor burden increases postoperatively indicating MRD. The patient had disease recurrence at 18 months. ROC, Receiver operating curve. MRD, minimal residual disease. SBRT, stereotactic body radiation therapy. ICI, immune checkpoint inhibition.

In the postoperative plasma samples, we defined MRD as a Z score in excess of the same 90% detection threshold previously defined in preoperative samples. MRD-EDGE detected postoperative MRD in 8 / 19 samples on plasma drawn a median of 43 days after surgery, four of which had confirmed disease recurrence. Postoperative MRD was found to be associated with shorter disease-free survival (**Fig 3c**) over a median follow-up of 49 months (range 18-76). Recurrence was not observed in any of the 11 patients in whom ctDNA was not detected. Of the 4 patients with postoperative detection who did not show evidence of recurrence, 1 received adjuvant therapy that may have eliminated residual disease, which has been demonstrated in other liquid biopsy settings^23^. One patient had short overall survival at 18 months (unrelated death), below the median time to recurrence in CRC^47^, and the remaining 2 patients had microsatellite unstable tumors that have been shown to be associated with prolonged time to relapse and occasional spontaneous regression^48,49^.

### Tracking of plasma tumor burden throughout neoadjuvant therapy with MRD-EDGE

We next sought to apply our MRD-EDGE SNV classifier to the challenging case of tracking plasma tumor burden in response to neoadjuvant immunotherapy. Tracking tumor burden in this setting could help optimize care during the crucial period between early-stage lung cancer detection and definitive surgery, with clinical implications such as extent of surgery planning for responders or moving to early surgery for non-responders. We evaluated plasma from three patients with early-stage NSCLC on a neoadjuvant immunotherapy protocol^50^ that randomized patients with early NSCLC to treatment with the ICI agent durvalumab with or without stereotactic body radiation therapy (SBRT) followed by surgical resection. Plasma was collected prior to the first ICI treatment or following day 3 SBRT (if applicable), at cycle 2 of ICI, prior to surgical resection, and after surgery (**Fig 3d**).

To determine an appropriate specificity threshold for use in neoadjuvant lung cancer monitoring, we applied MRD-EDGE to a cohort of early-stage LUAD patients evaluated previously^28^. MRD-EDGE maintained performance in this cohort compared to MRDetect (**Extended Data 4c-d**) and allowed us to identify a Z score detection threshold in a larger, orthogonal cohort.

We detected preoperative ctDNA in each of these three neoadjuvant treatment patients using the detection threshold prespecified from our early-stage LUAD cohort. One patient, Neo-01 (LUAD histology), had a marked decrease in plasma TF following SBRT, but ultimately plasma TF rose prior to surgery demonstrating a lack of response to ICI (**Fig 3f**). This patient had detectable ctDNA postoperatively and was found to have disease recurrence at 18 months following surgery. Two patients who did not receive SBRT showed minimally changed tumor burden throughout ICI treatment and no evidence of pathological response at the time of surgery. The first, Neo-02 (non-specific histology), had undetectable ctDNA postoperatively and remains free of disease at 29 months. The second, Neo-03 (squamous histology), was found to have postoperative MRD and recurred at 12 months after surgery (**Fig 3e**). These data highlight the potential of serial ctDNA monitoring during multi-pronged therapeutic regimens to define response to treatment and create opportunities for real-time therapeutic optimization.

### MRD-EDGE detects ctDNA shedding in precancerous adenomas and minimally invasive pT1 carcinomas

Whether noninvasive (precancerous) lesions shed ctDNA remains unresolved. The issue carries important implications for emerging early detection efforts where the presence of ctDNA from precancerous lesions may be advantageous in some settings, or alternatively diminish the precision of liquid biopsy screening tests. While MRD-EDGE requires a tumor prior and therefore cannot be used for screening, we reasoned that the exquisite sensitivity of our approach could nonetheless address whether ctDNA is shed from adenomas and polyp cancers (pT1pN0), where ctDNA detection through existing methods such as droplet digital PCR and targeted sequencing has been limited^51,52^.

We evaluated pre-resection plasma from 28 patients with malignant and premalignant lesions detected through screening at the Danish National Colorectal Screening Program^53^. Nine patients had pT1 lesions (defined as invasion of the submucosa but not the muscular layer, the earliest form of clinically relevant CRC^54^), and 19 patients had screen-detected precancerous adenomas (including one adenoma with microsatellite instability). As a positive control, we also evaluated plasma from 5 patients with metastatic CRC. We compared these samples to healthy control plasma that was sequenced at the same location and with the same platform as our adenoma and pT1 lesion plasma (‘Control Cohort B’, **Supplementary Table 1** and Methods). Consistent with prior reports^55–57^, we found decreased aneuploidy in adenomas (median 235Mb of genomewide aneuploidy) compared to our early-stage CRC samples (median 594Mb aneuploidy, *P*=0.02).

We next assessed performance of MRD-EDGE in this cohort. To ensure generalizability of detection, we applied the prespecified Z score threshold values from our preoperative early-stage CRC cohort (**Fig 3a-b**). These thresholds yielded similar specificity for adenoma and pT1 detections for both SNVs and CNVs (89% and 93%, respectively) in this separate cohort of control plasma samples sequenced with Illumina NovaSeq v1.5 rather than Illumina HiSeq X (**Supplementary Table 1**).

MRD-EDGE detected ctDNA shedding in 8 / 9 (89%) pT1 lesions and 8 / 19 (42%) precancerous adenomas (**Fig 4a**). Detection AUCs were higher for pT1 lesions than adenomas for both our SNV and CNV platforms, demonstrating decreased ctDNA signal in adenomas as expected (**Fig 4b**). As in our early-stage CRC cohort, we also analyzed performance in a cross-patient analysis (**Extended Data 5b-c**) and found similar detection ability. We further note that our patient-specific mutational compendium in this setting was drawn from formalin-fixed paraffin-embedded (FFPE) tissue samples, which are prone to more SNV artifacts^58^ than fresh frozen tissue samples used in our CRC and LUAD cohorts, further supporting the generalizability of classifiers among diverse tissue preparations. Using SNV-based TF estimations (Methods), we found lower TFs in detected lesions (median 2.88*10^-6^, range 1.02*10^-6^–1.45*10^-5^ in pT1 lesions and 3.78*10^-6^, range 1.17*10^-6^–1.21*10^-5^ in adenomas) than early-stage and metastatic CRC samples (**Fig 4c**). Detections for pT1 and adenoma lesions were significantly above our expected false positive rate of 10% (binomial *P*=2.1*10^-5^ and 2.1*10^-2^, respectively).

**Figure 4:**
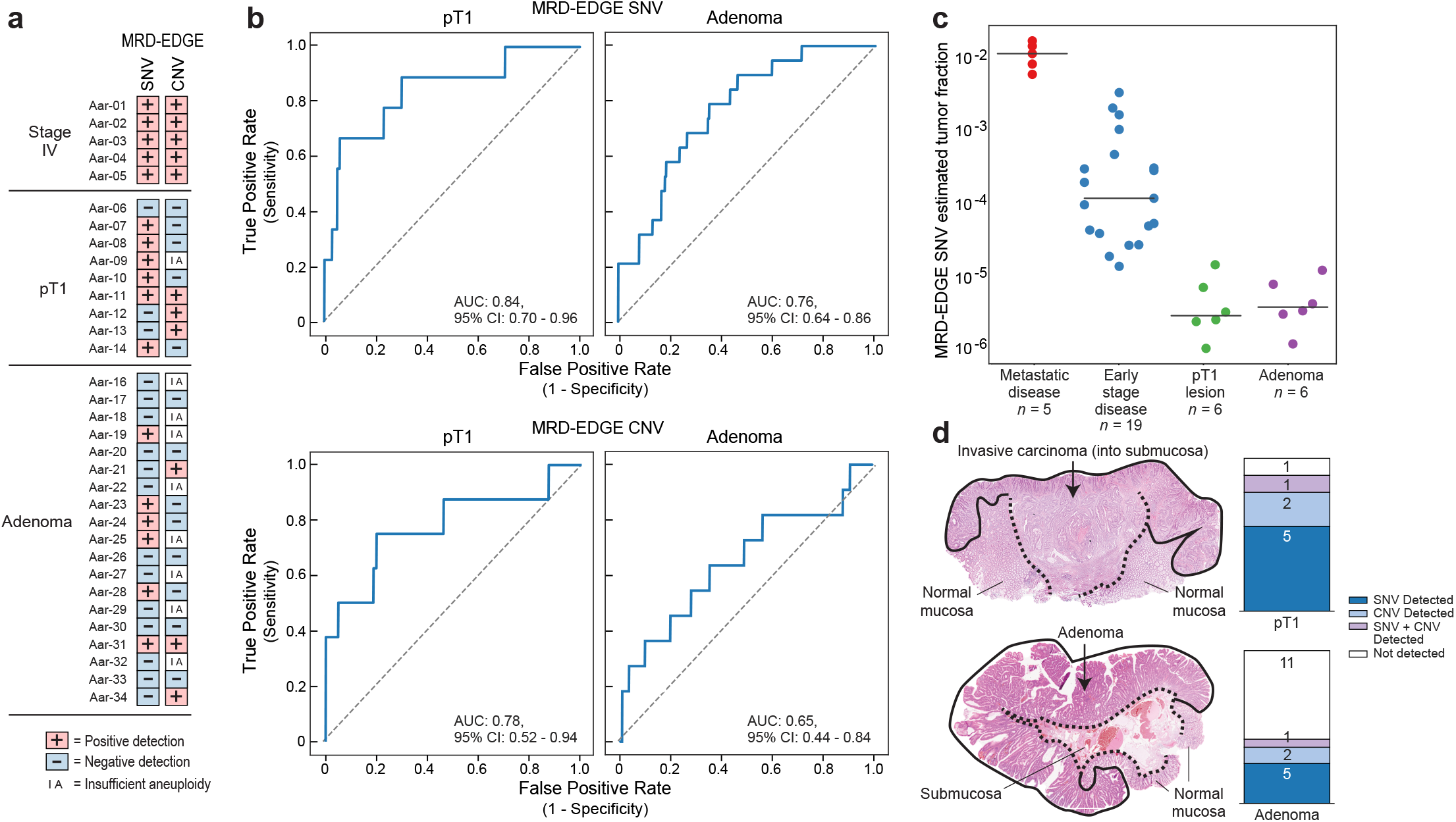
MRD-EDGE tumor-informed detection of ctDNA from screen-detected adenomas and pT1 lesions. **a)** Detection status of the cohort of Stage IV colorectal (CRC, *n*=5), screen-detected pT1 lesions (*n*=9) and screen-detected adenoma plasma samples (*n*=19) according to MRD-EDGE SNV and CNV classifiers. Samples with a Z score in excess of the detection threshold as prespecified in the early-stage CRC cohort (**Fig 3a-b**) are highlighted. **b)** ROC analysis for MRD-EDGE SNV (top) and CNV (bottom) classifiers in screen-detected adenomas (left) and pT1 lesions (right). Preoperative plasma samples were used as the true label, and the panel of control plasma samples (Control Cohort B) against all patient mutational compendia were used as the false label. For SNVs, 4 of 15 control samples were used in SNV model training and thus excluded from this analysis, yielding 11 control samples as a comparator. For CNVs, 5 of 15 control samples were used in a panel of normal samples (PON) for our read depth classifier (**Fig 2a**) and thus excluded from this analysis, yielding 10 control samples as a comparator. **c)** Plasma TF inference using genome-wide SNV integration for Stage IV CRC (*n*=5), early-stage preoperative CRC (*n*=19), SNV detected pT1 lesions (*n*=6), and SNV detected adenomas (*n*=6) shows decreasing estimated TF by CRC stage. Lines indicate median estimated TF. **d)** (left) histology image of the pT1 lesion Aar-14 (top) demonstrates invasion of the submusoca by dysplastic cancer cells, while an image of the adenoma Aar-17 (bottom) demonstrates the presence of dysplasia and absence of submucosal invasion. (right) barplots demonstrate number of plasma samples with detected ctDNA in patients with pT1 lesions (top) and adenomas (bottom). Detections are shaded by dark blue (MRD-EDGE SNV detections), light blue (MRD-EDGE CNV detections), light purple (SNV and CNV detections), and white (non-detected). ROC, receiver operating curve.

These data demonstrate that even without a significant invasive component, dysplastic tissue may shed ctDNA. The contribution of precancerous lesions or even benign clonal outgrowths to the cfDNA pool may thus form an important consideration as advanced non-tumor informed methods are deployed clinically, both for detection of adenomas and for early cancer detection efforts.

### MRD-EDGE enables ctDNA monitoring in melanoma plasma WGS without matched tumor

Across solid tumors, tumor tissue may be scarce due to considerations ranging from scant biopsy material (e.g., stage II melanoma), lack of primary biopsies at tertiary care centers, or restrictions on access to primary tissue. For example, in prior bespoke panel studies the requirement for matched tissue led to the exclusion of a substantive proportion of eligible patients due to low tumor DNA purity or quality^20,59^. Further, in several cancers, non-surgical treatment modalities like radiation are given with curative intent, again limiting opportunities for tumor-informed approaches. This introduces the need for tumor-agnostic (*de novo*) mutation calling platforms for clinical surveillance. Our improved signal to noise enrichment in the tumor-informed setting (**Extended Data 1d**) motivated us to consider *de novo* mutation calling using our MRD-EDGE platform. In this setting, there is no *a priori* knowledge of high likelihood mutated loci, and ctDNA signal is therefore far more challenging to distinguish from sequencing error.

*De novo* mutation calling with MRD-EDGE requires the evaluation of all plasma fragments that harbor SNVs, which range from 1*10^7^-1*10^8^ per plasma sample in our WGS cohorts (Methods, **Supplementary Table 1**). As these SNVs harbor far greater cfDNA sequencing noise compared to ctDNA signal, we reasoned that higher specificity thresholds would need to be applied to the output of the deep learning classifier. To determine an appropriate *de novo* specificity threshold for our MRD-EDGE deep learning SNV classifier (**Fig 1d**) we used the same *in silico* admixtures as in the tumor-informed setting (validation melanoma sample MEL-01 admixed with a held-out healthy control plasma sample, **Fig 1e**). We compared signal to noise enrichment with detection AUC at different specificity thresholds imposed on the MRD-EDGE ensemble model output (**Extended Data 6a-b**, Methods) to find an optimal threshold for classification of ultrasensitive TFs (TF 5*10^-5^). As expected, our empirically chosen threshold in the *de novo* classification context (0.995) was higher than the balanced threshold (0.5) used in the tumor-informed setting. At this threshold, AUC for ultrasensitive detection (5*10^-5^) was 0.77 (**Fig 5a**). Signal to noise enrichment for MRD-EDGE was 2,518 fold (range 1,817-3,058 fold) compared to the MRDetect SVM (mean 8.3 fold, range 8-9 fold) in a matched analysis performed with the same samples used in the tumor-informed setting (**Extended Data 1d**). This equates to 301-fold (range 211-357 fold, **Fig 5b**) higher enrichment for MRD-EDGE compared to MRDetect.

**Figure 5:**
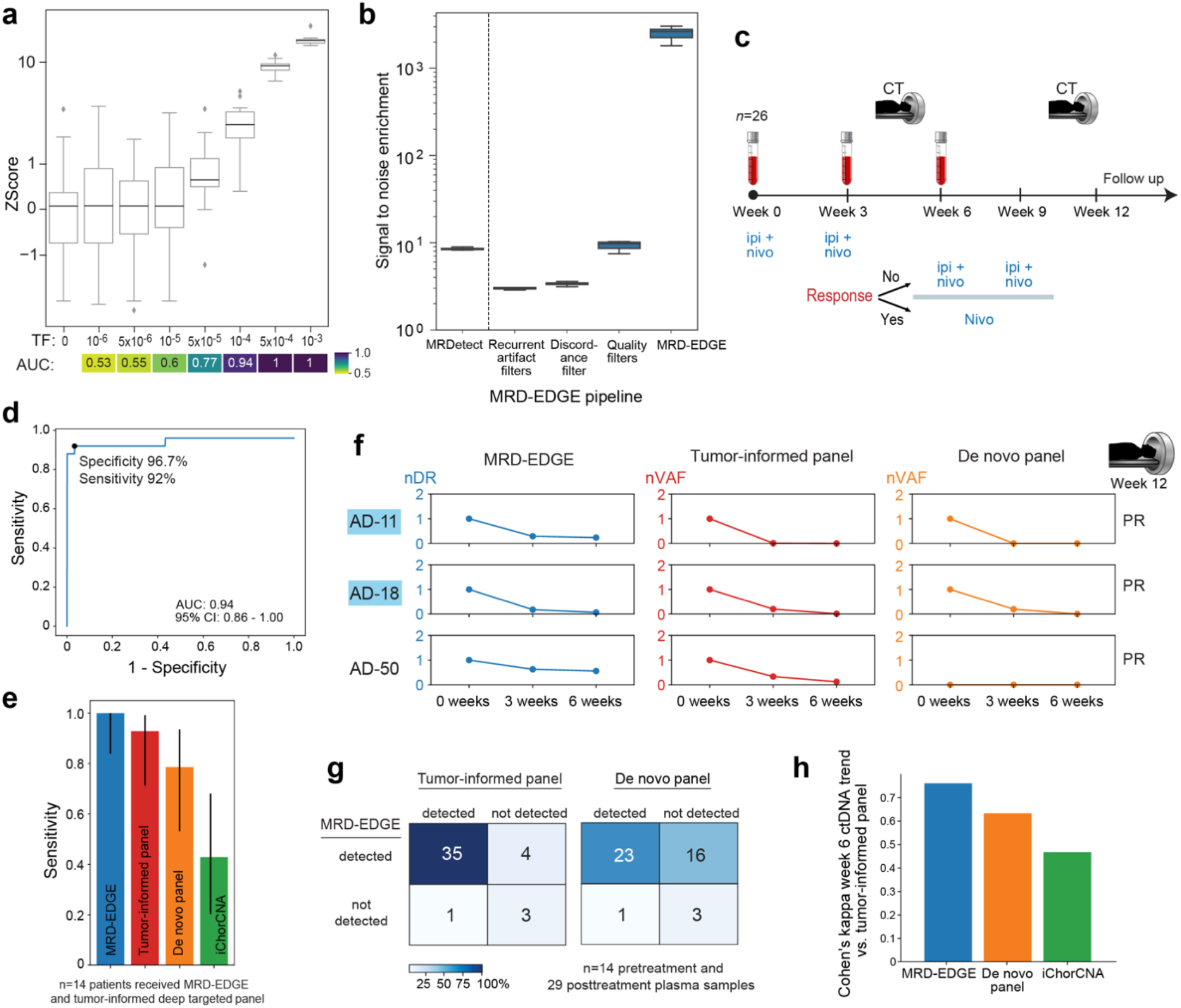
MRD-EDGE accurately monitors ctDNA in melanoma plasma WGS without matched tumor. **a)** *In silico* studies of cfDNA from the metastatic melanoma sample MEL-01 (pretreatment TF of 3.5%) mixed in *n*=20 replicates against cfDNA from a healthy plasma sample (TF=0) at mix fractions 10^-6^–10^-3^ at 16X coverage depth. MRD-EDGE enables sensitive TF detection as measured by Z score against healthy controls at TF=5*10^-5^ (AUC 0.77) without matched tumor tissue to guide SNV identification. Box plots represent median, bottom and upper quartiles; whiskers correspond to 1.5 x IQR. An AUC heatmap measures detection vs. TF=0 at different mixed TFs. **b)** Signal to noise enrichment analysis for MRDetect SVM and for each step of the MRD-EDGE *de novo* mutation calling pipeline. Final pipeline enrichment is 2,518-fold for MRD-EDGE vs. 8.3-fold for the MRDetect SVM in the same plasma samples. MRD-EDGE provides for a cumulative 301-fold enrichment over MRDetect. **c)** Study schematic for adaptive dosing melanoma cohort (*n*=26 patients with advanced melanoma). All patients began treatment with combination ipilimumab (3 mg/kg) and nivolumab (1 mg/kg). Plasma was collected at pretreatment timepoint at week 0, at second dose of combination ICI at Week 3, and at Week 6. Beginning at Week 6 patients received either combination ICI or ICI monotherapy based on imaging response: patients with stable or shrinking disease on Week 6 CT received nivolumab monotherapy and those with tumor growth received additional combination therapy. Further CT imaging was performed at Week 12. **d)** ROC analysis for the detection of pretreatment melanoma using MRD-EDGE for healthy individuals (*n*=30, false label) and patients with melanoma (*n*=25, true label). One pretreatment melanoma plasma sample with high TF used in model training was withheld from this analysis. Detection rate cutoff was selected as the first operational point with specificity of 90% or greater. **e)** Fourteen of 26 patients from the adaptive dosing cohort underwent sequencing with a tumor-informed targeted panel^8^ (‘tumor-informed panel’). Vertical bars demonstrate pretreatment detection sensitivity for MRD-EDGE, the tumor-informed panel, a *de novo* panel based on the *de novo* calling thresholds^8^ used for the tumor-informed panel, and ichorCNA. Error bars represent 95% binomial confidence interval for empiric sensitivity within 14 trials. **f)** Serial tumor burden monitoring on ICI with MRD-EDGE, tumor-informed panel, and *de novo* panel for 3 patients with melanoma. Tumor burden estimates are measured as a detection rate normalized to the pretreatment sample (normalized detection rate, nDR) for MRD-EDGE and as variant allele fraction (VAF) normalized to the pretreatment VAF (normalized VAF, nVAF) in the tumor-informed panel and *de novo* panel. MRDetect accurately captures trends in TF, while the *de novo* panel faces sensitivity barriers in low TF settings where plasma VAF < 0.005. Blue highlights surrounding sample name indicate samples with 14 or more SNVs covered in the tumor-informed panel. **g)** Forty-three pre- and posttreatment samples from the adaptive dosing melanoma cohort underwent sequencing with MRD-EDGE and the tumor-informed panel. (top) Heatmap demonstrating detection overlap (measured as the agreement between platforms of detected ctDNA and undetectable ctDNA) between MRD-EDGE and the tumor-informed panel shows high concordance (88%) between the two platforms. (bottom) Lower detection overlap (60%) is present between MRD-EDGE and the *de novo* targeted panel due to sensitivity floors in the *de novo* panel. **h)** Barplot of Cohen’s kappa agreement metric for Week 6 ctDNA trend (increase or decrease) compared to pretreatment baseline between 3 mutation callers and the tumor-informed panel: MRD-EDGE, *de novo* panel, and iChorCNA. MRD-EDGE demonstrates most agreement with the tumor-informed panel (Cohen’s Kappa 0.75). ROC, Receiver operating curve. IQR, interquartile range. IQR, interquartile range. CT, computed tomography.

After benchmarking fragment-level performance for *de novo* mutation calling with MRD-EDGE, we evaluated performance at the sample level in a cohort of patients with advanced cutaneous melanoma treated with combination ICI on The Adaptively Dosed Immunotherapy Trial^60^ (‘adaptive dosing cohort’, *n*=26 patients, 2-4 timepoints per patient, **Fig 5c**). In this cohort, plasma was sampled at baseline (pretreatment) and prior to the second (Week 3) and third (Week 6) infusion of the ICI agents nivolumab and ipilimumab. The protocol aimed to spare excess combination ICI treatment by identifying responders through early imaging at Week 6 and transitioning these patients to monotherapy with nivolumab.

We compared ctDNA detection rates in our melanoma cohort to a cohort of controls (*n*=30 patients without known cancer, ‘Control Cohort C’) sequenced under similar conditions (Illumina NovaSeq v1.0 for melanoma and control groups) to avoid inter-platform bias. MRD-EDGE identified ctDNA in pretreatment plasma from cutaneous melanoma samples (*n*=25 after holding out one melanoma plasma sample with high TF used in neural network training), yielding an AUC of 0.94 (95% CI: 0.86-1.0, **Fig 5d**). In keeping with our tumor-informed analyses, we chose the first detection threshold at a specificity of 90% or greater (sensitivity of 92%, specificity of 96.7%). As a negative control, we included pre- and posttreatment plasma samples from a patient with acral melanoma (*n*=3 total plasma samples) within the same sequencing batch. As expected, we observed no ctDNA detection in these samples (**Extended Data 6c**), confirming that our classifier is specific for the distinct mutational signatures of cutaneous melanoma.

To benchmark MRD-EDGE ctDNA detection in pretreatment plasma against alternative methods, we compared results to a state-of-the-art targeted panel^8^ with tumor-informed mutation calling covering 129 common cancer genes (‘tumor-informed panel’) in a subset of 14 patients. Tumor-informed detection was based on an average of 9.4 panel-covered SNVs per sample (range 2-29, **Supplementary Table 4**). Four patients had 14 or more SNVs (highlighted in **Fig 5f**, **Extended Data 7**), a range comparable to leading bespoke panels^19,20,59^. In parallel, results were also compared to the same targeted panel with *de novo* mutation calling (‘*de novo* panel’) and to iChorCNA^10^, an established WGS CNV TF estimator. In cutaneous melanoma pretreatment plasma samples profiled across methods, sensitivity for MRD-EDGE ctDNA detection was 100% (binomial 95% CI 83.8%-100%), compared to 93% (71.2%-99.2%) for the tumor-informed panel, 79% (53.1%-93.6%) for the *de novo* panel and 43% for iChorCNA (20.2%-68.0%) (**Fig 5e**).

We next assessed MRD-EDGE’s ability to monitor changes in ctDNA TF following ICI treatment compared to alternative methods. Given the unknown variable of tumor mutational burden in these samples and the influence of mutation load on detection rate, MRD-EDGE trends in TF were measured as a detection rate normalized to pretreatment TF (‘normalized detection rate’, nDR). For comparison in targeted panels, VAF was normalized to the pretreatment timepoint (‘normalized VAF’, nVAF). Side-by side comparisons demonstrate broadly similar trends in tumor burden following ICI treatment (**Fig 5f, Extended Data 7)**.

We considered a sample detected by the tumor-informed panel if estimated VAF across all surveyed genes was greater than zero, while detection in the *de novo* panel was measured as variant allele frequency (VAF) > 0.005 per published methods^8^. Among samples evaluated across platforms (*n*=43 total, 14 pretreatment and 29 posttreatment samples), detection consistency (measured as the agreement between platforms of detected ctDNA and undetectable ctDNA) was highest between MRD-EDGE and the tumor-informed panel at 38 of 43 samples (88%, **Fig 5g** left). MRD-EDGE detected the lowest VAF detected by the tumor-informed panel, estimated at 1*10^-4^, validating our *in silico* benchmarking of detection sensitivity in clinical practice. Detection consistency was lower at 26 of 43 samples (60%) between MRD-EDGE and the *de novo* panel, likely due to the sensitivity floor of 0.005 in the latter method (**Fig 5g**, right). To benchmark MRD-EDGE’s utility in clinical surveillance, we compared changes in ctDNA TF at Week 6 following ICI treatment. Changes in nDR or nVAF showed higher agreement between MRD-EDGE and the tumor-informed panel, compared to the agreement with the *de novo* panel and iChorCNA (**Fig 5h**). In summary, MRD-EDGE enables ultrasensitive melanoma ctDNA detection and TF monitoring on par with an established tumor-informed panel.

### MRD-EDGE sensitively tracks response to immunotherapy in metastatic melanoma

In advanced melanoma, radiographic response may not be apparent for months after ICI initiation due to pseudo-progression or residual fibrous tissue^61,62^, limiting the sensitivity of imaging to detect meaningful changes in tumor burden. Further, the absence of biomarkers that predict which patients will respond to therapy can lead to excess or futile treatment in unselected populations^63^. Liquid biopsy can improve ICI care by providing faster readouts of response, orthogonal measurement of TF trends, and longitudinal noninvasive TF surveillance. Several panel approaches have demonstrated that changes in plasma TF as measured through increasing or decreasing ctDNA TF can complement imaging to predict response to ICI therapy^20,21,59,64,65^.

We sought to explore the clinical utility of *de novo* (i.e., non tumor-informed) MRD-EDGE in ICI-treated patients with metastatic melanoma. We expanded the adaptive dosing melanoma^60^ cohort described above (*n*=26 patients, **Fig 6a** right panel) to include additional patients treated with standard of care immunotherapy (‘conventional immunotherapy’, *n*=11 patients, **Fig 6a** left panel, Supplemental Table 4). As further demonstration of applicability across platforms, the adaptive dosing cohort was sequenced on Illumina NovaSeq v1.0 while the standard of care immunotherapy cohort was sequenced on Illumina HiSeq X (Supplemental Table 3). No tumor or matched normal tissue was used in this *de novo* plasma WGS analysis.

**Figure 6:**
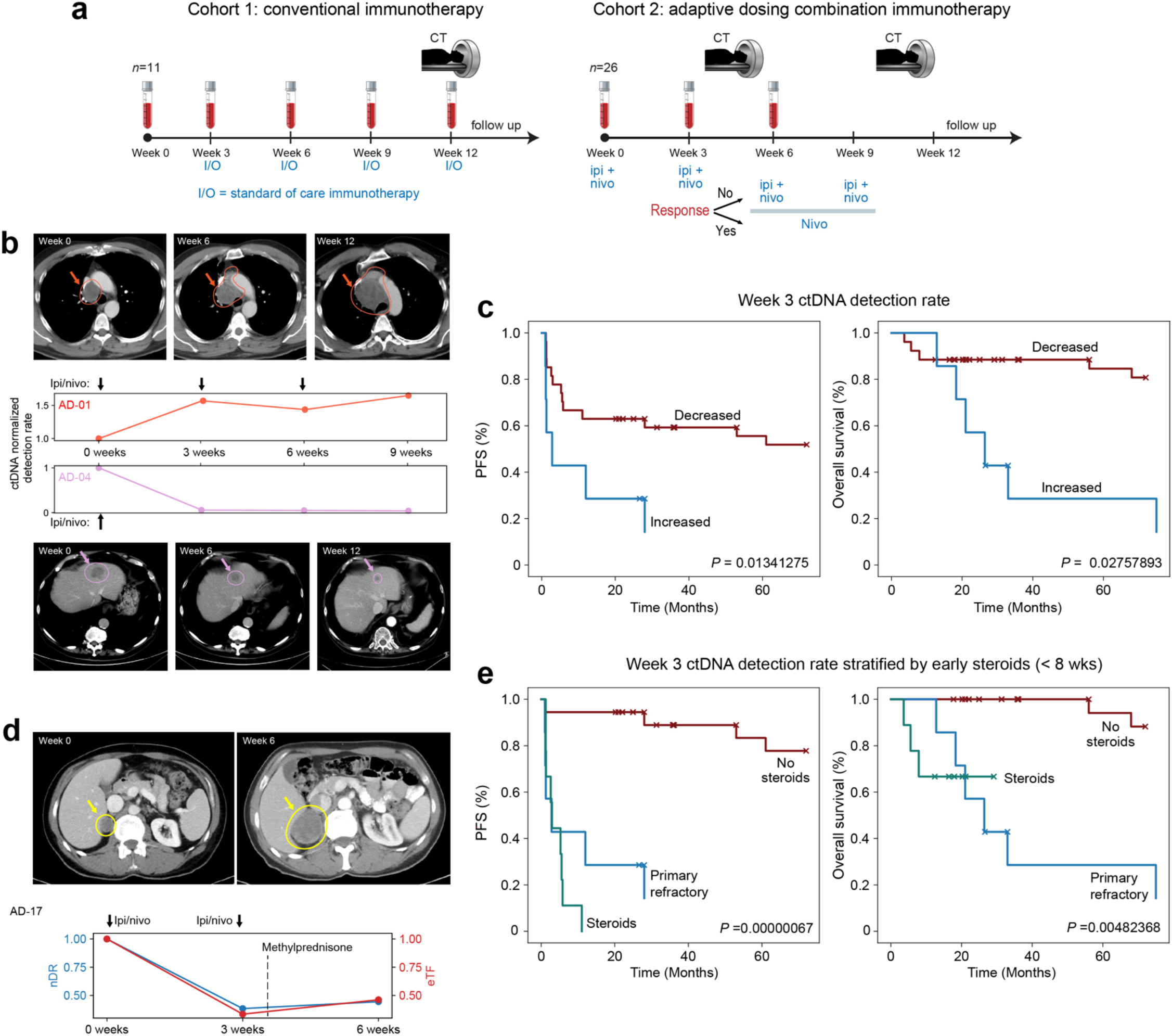
Serial monitoring of clinical response to immunotherapy with MRD-EDGE. **a)** Study schematics of two advanced melanoma cohorts. (left) conventional immunotherapy cohort received nivolumab monotherapy or combination ICI. Plasma was collected at pretreatment timepoint and weeks 3, 6, and 12. Cross sectional imaging to evaluate response to treatment was performed at 12 weeks. (right) adaptive dosing cohort received combination immunotherapy as described in **Fig 5c**. **b)** Serial plasma TF monitoring with MRD-EDGE corresponds to changes seen on imaging. TF estimates are measured as a detection rate normalized to the pretreatment sample (normalized detection rate, nDR) for MRD-EDGE. (top) ctDNA nDR grossly increases over time in a patient with disease refractory to ICI. The patient had progressive disease at Week 6 and Week 12 CT assessment. (bottom) ctDNA nDR decreased at Week 3 in a patient with a partial response to therapy. CT imaging demonstrates tumor shrinkage at Week 6 and Week 12. **c)** Kaplan-Meier progression-free and overall survival analysis for Week 3 ctDNA trend in patients with decreased (*n*=27) or increased (*n*=7) nDR as measured by MRD-EDGE. Patients with undetectable pretreatment ctDNA (*n*=3) were excluded from the analysis. Increased nDR at Week 3 shows association with shorter progression-free and overall survival (two-sided log-rank test). **d)** (top left) pretreatment CT imaging of a patient with decreased ctDNA in response to ICI at Week 3 on both MRD-EDGE (nDR, blue) and a tumor-informed panel (normalized variant allele frequency, nVAF, red). Following the administration of methylprednisone at Week 3, estimated TF on both ctDNA detection platforms increased. At Week 6, progressive disease is seen on CT imaging (top right). **e)** Early steroids for irAEs within the combination ICI dosing period (prior to Week 8) further stratify Week 3 survival analyses. Kaplan-Meier progression-free and overall survival analysis was performed on patients with primary refractory disease (‘primary refractory’, blue, *n*=7), defined as rising nDR seen at Week 3 following first dose of treatment, decreasing ctDNA who did not receive steroids (“no steroids”, red, *n*=18), and patients who received steroids for immune-related adverse events within the combination ICI dosing period (‘steroids’, green, *n*=9). P value reflects multivariate logrank test. ICI, immune checkpoint inhibition. CT, computed tomography.

Trends in MRD-EDGE nDR tracked radiographic imaging results. For example, in a patient who progressed on treatment, progressive disease was seen on computed tomography (CT) at Week 6 and Week 12 while nDR concomitantly increased (**Fig 6b**, top). Similarly, radiographic imaging demonstrated ongoing tumor shrinkage in a patient who responded to treatment, matched by a rapid and persistent decrease in nDR that occurred by Week 3 (**Fig 6b**, bottom).

We next evaluated MRD-EDGE’s ability to prognosticate clinical outcomes at serial plasma timepoints (122 pre- and posttreatment plasma samples from *n*=37 patients, **Supplementary Table 4**). Patients with undetectable pretreatment ctDNA (*n*=3) were excluded from further clinical analyses. We found that change in ctDNA nDR as measured by increased or decreased plasma TF following treatment to be predictive of both PFS (*P*=0.01) and OS (*P*=0.03, **Fig 6d**) as early as Week 3 after the first ICI infusion. This prognostic role for plasma TF changes after first ICI infusion and prior to any conventional imaging has also been noted in response to single-agent ICI in NSCLC^21^, and demonstrate a role for liquid biopsy TF surveillance in the earliest days of ICI treatment. We also found significant PFS and OS relationships for change in ctDNA nDR at Week 6 (**Extended Data 8a**). In contrast, CT imaging was available for our adaptive dosing cohort at Week 6, and here we found no significant relationship between RECIST response and OS (*P*=0.40, **Extended Data 8b**).

Notably, the first OS event in our Week 3 and Week 6 ctDNA survival analysis occurred in a patient with decreasing nDR at Week 3 and Week 6 who enrolled on protocol following prior treatment of brain metastases. CT imaging (partial response) and ctDNA trends for both MRD-EDGE and the tumor-informed panel identified an extracranial response to therapy. This patient, however, had intracranial progression at 5 months and was taken off protocol. Such findings are consistent with the melanoma ctDNA literature, where ctDNA trends are known to reflect extracranial rather than intracranial tumor burden^66^, and suggest that ctDNA monitoring should be used with caution in patients at high risk of intracranial progression.

Despite significant PFS and OS relationships for ctDNA trends at Week 3, we noted several instances in which decreasing Week 3 nDR was not indicative of durable ICI response. We reasoned that the high toxicity rate from combination ICI, where nearly 40% of patients will stop treatment early because of immune-related adverse events (irAEs)^67^, may have confounded classification at Week 3. Clinically, severe irAEs are often treated with corticosteroids, and early steroid use (within 8 weeks of ICI treatment) is associated with shorter PFS and OS in melanoma^68^. We therefore stratified our melanoma patients into 3 groups, patients with primary refractory disease (initial increase in ctDNA nDR, *n*=7), and patients with an initial ctDNA response either treated or untreated with early steroids (*n*=9 and *n*=18, respectively). This classification proved strongly predictive of both PFS (*P*=1.3*10^-7^) and OS (*P*=1.7*10^-4^, **Fig 5f**), and suggests that early treatment responses, measured via ctDNA, may be inhibited by steroids. In summary, with no need for matched tumor and a standard WGS workflow, MRD-EDGE offers the potential for real-time serial monitoring of plasma ctDNA in conjunction with imaging to assess immunotherapy response.

## DISCUSSION

The use of noninvasive liquid biopsy to detect MRD and track response to therapy heralds the next frontier in precision oncology. We previously observed that the sensitivity of deep targeted sequencing approaches may be limited in the context of low plasma TF (e.g., MRD or the nadir of response to immunotherapy), and used WGS of plasma to expand the number of informative sites and therefore increase sensitivity in this setting. Herein, we introduced a machine learning-based classifier MRD-EDGE that is designed to integrate an expanded feature set for SNVs and CNVs to substantially enhance ctDNA signal enrichment.

Broadly, MRD-EDGE can leverage both prior knowledge of tumor-specific mutational compendia and a biologically-informed feature space to enrich ctDNA signal. Our MRD-EDGE SNV deep learning strategy differs markedly from other deep learning variant callers^69,70^ through our use of disease-specific biology to inform somatic mutation identification. Our focus on classifying fragments rather than loci allowed us to overcome the inability to apply consensus mutation calling, the cornerstone of most variant calling strategies, in extremely low TF settings. Moreover, fragment-based classification enabled an increase in the size of our training corpuses to hundreds of thousands of observations, which is critical to comprehensive pattern recognition with neural networks^71^. The deep learning SNV architecture in MRD-EDGE provides a flexible platform for integrating disease-specific molecular features, outperforms other machine learning approaches, and demonstrates generalizability across cancer types and sequencing preparations.

For CNVs, machine-learning guided signal denoising enables accurate inference of plasma read depth skews, while fragmentomics and BAF provide orthogonal metrics for CNV assessment. Our use of tumor-specific copy number profiles combined with powerful denoising enables increased sensitivity compared to established read depth approaches^10,11^. The use of neutral segments as a sample level internal control offers an additional specificity advantage compared to tumor-agnostic fragment-based methods^9,23^. The lower degree of aneuploidy needed for ultrasensitive detection (**Fig 2e**) and ability to capture signal from copy-neutral LOH will enable application to a diverse set of solid tumors even in the absence of high somatic SNV burden (**Extended Data 3**).

We expect that the simple WGS workflow, which obviates the need for custom panel generation and molecular barcodes, and ability to work with limited input material (1 mL of plasma) will enhance MRD-EDGE translational impact in diverse clinical settings, especially given the rapid decline in raw sequencing costs. MRD-EDGE enabled the detection of postoperative CRC and LUAD MRD, as well as tracking of plasma TF dynamics in response to neoadjuvant ICI. Our data highlight the potential for real-time therapeutic optimization in the neoadjuvant setting, which could potentially inform early surgery or treatment change for non-responders in order to maximize curative opportunity.

The distinct sensitivity of MRD-EDGE allowed us to examine the detection of ctDNA shedding from precancerous colorectal adenomas. While this tumor-informed approach cannot be used for screening, the detection of ctDNA in a substantial proportion of cased argues that ctDNA may be present without invasive disease. This carries important implications for ongoing efforts to develop liquid biopsy approaches for cancer screening^9,13,72,73^. Considering the value of precancerous lesion detection in CRC screening^74^, these data demonstrate that ctDNA-guided detection of premalignant lesions is a viable goal, provided that tools with sufficient sensitivity can be developed for this setting. On the other hand, the demonstration of ctDNA shedding without an invasive component suggests that clonal mosaicisms in normal tissues may impact cancer screening efforts in a manner similar to the observation of confounding clonal hematopoiesis mutations in targeted sequencing^73,75–77^. This may be particularly important for hotspot mutations given the pervasive nature of clonal outgrowths^78–80^ and the potential of the plasma to aggerate signal across potentially thousands of separate clones. Similarly, it is unknown to what degree normal solid tissue clonal outgrowths differ from malignant counterparts in fragment length or methylation profiles, which may impact non-mutational ctDNA screening methods.

We further leveraged the enhanced signal to noise enrichment of MRD-EDGE to perform *de novo* (non-tumor informed) SNV mutation detection in advanced melanoma. The emerging role of early ctDNA trends in monitoring ICI response, seen here and elsewhere^20,21,59^, is reflected in the recent Center for Medicare & Medicaid Services approval of tumor-informed bespoke assays to prognosticate response to immunotherapy after 6 weeks. In the phase 2 trial^20^ that led to this approval, the requirement for a matched tumor sample for bespoke panel design led to the exclusion of one-third of patients due to low tumor DNA purity or quality. In contrast, MRD-EDGE required only plasma, and produced performance on par with a comparable tumor-informed panel. MRD-EDGE allowed for early and accurate assessment of response to ICI, a challenging clinical setting for prognostication^63,64^. Future large-scale interventional studies will be critical to demonstrate the value of rapid and quantitative estimation of ICI response to inform real-time clinical decision making.

Collectively, our data support the use of plasma WGS as a complementary strategy to the prevailing paradigm of ctDNA mutation detection via deep targeted panel sequencing. Our approach can complement targeted panels, as well as other liquid biopsy tools such as methylation-based assays, to create a comprehensive liquid biopsy toolkit that tailors sequencing approach to clinical application. For example, we envision that improved cancer screening through early detection efforts will allow the diagnosis of cancers at less advanced stages^9,12,13,73^. Low tumor-burden disease treated with surgical and/or non-surgical means will benefit from ultra-sensitive TF monitoring via MRD-EDGE. In the event of high burden disease relapse, deep targeted panels^5,6,8,19,21^, better suited to provide mutational profiling through exhaustive coverage depth, can nominate gene targets for systemic targeted therapy. While the value of therapy optimization based on MRD-EGDE monitoring requires investigation in large clinical cohorts, our findings highlight the potential of ctDNA as a quantitative tumor burden biomarker that provides real-time feedback in response to therapy and early insight into relapsed disease.

## ACKNOWLEDGEMENTS

We extend our thanks to the patients who contributed plasma and tissue to this project as well as their families. We thank and acknowledge the Danish Cancer Biobank and the Endoscopy III study team (Lars Nannestad Jørgensen and Morten Rasmussen (Bispebjerg Hospital, Copenhagen), Mogens R. Madsen and Anders H. Madsen (Herning Hospital, Herning), Linnea Ferm and Eva Rømer (Hvidovre Hospital, Hvidovre), Tobias Boest and Berit Andersen (Randers Hospital, Randers) and Ali Khalid (Viborg Hospital, Viborg) for providing access to blood and tissue materials. We also thank the Landau laboratory and the NYGC computational biology and sequencing teams for help and feedback throughout this work. This work was supported by the Mark foundation Aspire Award (DAL and CLA); Novo Nordisk Foundation [grant number NNF170C0025052 (CLA)]; the Danish Cancer Society [grant numbers R146-A9466-16-S2 (CLA); R231-A13845 (CLA); and R257-A14700 (CLA). MSKCC investigators are supported by Cancer Center Support Grant P30 CA08748 from the National Institutes of Health/National Cancer Institute. AJW received support from the Conquer Cancer Foundation Young Investigator Award. DAL is supported by the Burroughs Wellcome Fund Career Award for Medical Scientists and the National Institutes of Health (NIH) Director’s New Innovator Award (DP2-CA239065). The opinions, results, and conclusions reported in this paper are those of the authors and are independent from these funding sources.

## METHODS

### Human subjects and sample processing

This study was approved by the local ethics committee and by the institutional review board (IRB) and was conducted in accordance with the Declaration of Helsinki protocol. Blood samples were collected from patient and healthy adult volunteers enrolled in clinical research protocols at NewYork-Presbyterian/Weill Cornell Medical Center, Memorial Sloan Kettering Cancer Center, Massachusetts General Hospital, the Royal Marsden NHS Foundation Trust in the United Kingdom, or Aarhus University Hospital, Bispebjerg Hospital, Randers Hospital, Herning Hospital, Hvidovre Hospital, and Viborg Hospital in Denmark. Melanoma tumor, normal and plasma samples from the Royal Marsden NHS Foundation Trust were obtained under an ethically approved protocol (Melanoma TRACERx, Research Ethics Committee Reference 11/L0/0003). Tumor tissues were collected from resected lung, melanoma, colorectal cancer, and adenoma specimens. The diagnosis of cutaneous melanoma, NSCLC, CRC, and adenoma was established according to World Health Organization criteria and confirmed in all cases by an independent pathology review. Informed consent on IRB-approved protocols for genomic sequencing of patients’ samples was obtained before the initiation of sequencing studies.

### Germline and tumor DNA processing

Tumor tissue and matched germline DNA from peripheral blood mononuclear cells (PBMCs) or adjacent normal tissue were collected and stored at −80 °C until they were processed for extraction. Genomic DNA was extracted from tumor tissue using the QIAamp DNA Mini Kit (Qiagen). Genomic DNA was extracted from PBMCs using the QIAamp DNA Blood Kit (Qiagen). Libraries were prepared using the TruSeq DNA PCR-Free Library Preparation Kit (Illumina) with 1 μg of DNA input after the recommended protocol^84^, with minor modifications as described below. Intact genomic DNA was concentration normalized and sheared using the Covaris LE220 sonicator to a target size of 450 bp. After cleanup and end repair, an additional double-sided bead-based size selection was added to produce sequencing libraries with highly consistent insert sizes. This was followed by A-tailing, ligation of Illumina DNA Adapter Plate adapters and two post-ligation bead-based library cleanups. These stringent cleanups resulted in a narrow library size distribution and the removal of remaining unligated adapters. Final libraries were run on a Fragment Analyzer (Agilent) to assess their size distribution and quantified by qPCR with adapter-specific primers (Kapa Biosystems). Libraries were pooled together based on expected final coverage and sequenced across multiple flow cell lanes to reduce the effect of lane-to-lane variations in yield. WGS was performed on the HiSeq X (HCS HD 3.5.0.7; RTA v2.7.7) or NovaSeq v1.0 (Illumina) at 2 x 150-bp read length, using SBS v3 (**Supplementary Table 1**).

Plasma DNA processing. At the same day of blood collection, blood collection tubes (Streck or K2-EDTA, **Supplementary Table 1**) were centrifuged at 2,000 r.p.m. for 10 min to separate plasma. cfDNA was then extracted from human blood plasma by using the Mag-Bind cfDNA Kit (Omega Bio-Tek). The protocol was optimized and modified to optimize yield^28^. Elution time was increased to 20 min on a thermomixer at 1,600 r.p.m. at room temperature and eluted in 35-μl elution buffer. The concentration of the samples was quantified by a Qubit Fluorometer (Thermo Fisher), and samples were run on a fragment analyzer by using the High Sensitivity NGS Fragment **Analysis Kit (Agilent) to define the size of cfDNA extracted and genomic DNA contamination.** For plasma samples that were found to have significant genomic DNA contamination (fragment size > 240 base pairs for more than 20% of fragments at library preparation) we performed a 0.4x cleanup using SPRIselect magnetic beads (Beckman Coulter) on the extracted cfDNA.

A subset of plasma samples was sequenced at Aarhus University in Denmark (**Supplementary Table 1**). For these samples, blood samples were collected in K2-EDTA 10 mL tubes (Becton Dickinson). Within two hours of blood collection, the tubes were centrifuged at 2,000 r.p.m. for 10 minutes to separate plasma. Isolated plasma was centrifuged again at 2,000 r.p.m. for 10 minutes. cfDNA was then extracted from human blood plasma using the QIAmp Circulating Nucleic Acids kit (Qiagen), eluted in 60-μl elution buffer (10 mM Tris-Cl, pH 8.5). The concentration of the samples was quantified by droplet digital PCR (ddPCR, Bio-Rad Laboratories), using assays specific to two highly conserved regions on Chr3 and Chr7, as previously described^85^. In addition, all samples were screened for contamination of genomic DNA from leucocytes using a ddPCR assay targeting the VDJ rearranged IGH locus specific for B cells, as previously described^85^. No samples were contaminated by genomic DNA from leucocytes.

### Plasma cfDNA library preparation and sequencing

Samples sequenced at the New York genome Center were processed using KAPA Hyper Library Preparation. Cohorts included in Zviran et al. were processed as previously described^28^. Samples with a mass above 5 ng were prepared for next-generation sequencing on Illumina’s HiSeq X or NovaSeq by using a modified manufacturer’s protocol. The protocol was scaled down to half reaction by using 25μl of extracted cfDNA. IDT for Illumina TruSeq Unique Dual Indexes^84^ was used by diluting 1:15 with EB (elution buffer), and ligation reaction was adjusted to 30 minutes. Additional 0.8x SPRIselect magnetic beads (Beckman Coulter) cleanup was included after post-ligation cleanup to remove excess adapters and adapter dimers. cfDNA from 1 mL of plasma was used for all of the plasma samples in this study. For samples with low concentration, an additional 1 ml of plasma was extracted, and the DNA aliquot with the highest mass was used for library preparation. The number of PCR cycles was dependent on initial cfDNA total mass. For samples with more than 5 ng of total cfDNA, 5-7 PCR cycles were performed. For samples with less than 5 ng of total cfDNA, 7-10 PCR cycles were performed (**Supplementary Table 1**). Quality metrics were performed on the libraries by Qubit Fluorometer, High Sensitivity DNA Analysis Kit and KAPA SYBR FAST qPCR Kit (Roche). WGS was performed on the HiSeq X (HCS HD 3.5.0.7; RTA v2.7.7) at 2 × 150-bp read length or NovaSeq v1.0 at 2 x 150-bp read length (**Supplementary Table 1**) to a target depth of 30x.

Plasma samples sequenced at Aarhus University also used KAPA Hyper Library Preparation. cfDNA from 2mL plasma (see **Supplementary Table 1** for DNA mass) was used as input for library preparation using a modified manufacturer’s protocol. xGen UDI-UMI Adapters were used and the ligation reaction was adjusted to 30 minutes. Agencourt AMPure XP beads (Beckman Coulter) were used for both cleanup steps with a bead:DNA ratio of 1.2x and 1.0x for the post-ligation and post-PCR cleanup, respectively. The number of PCR cycles was 7 for all cfDNA samples. Qubit Fluorometer and TapeStation D1000 were used for library quality control. WGS was performed on NovaSeq v1.5 at 2 x 150-bp read length to a target depth of 30x.

### Preprocessing, quality control analysis and sample identification and concordance

WGS reads for primary tumor, matched germline and plasma samples were demultiplexed using Illumina’s bcl2fastq (v2.17.1.14) to generate FASTQ files. The primary tumor and matched germline WGS were submitted to the New York Genome Center somatic preprocessing pipeline, which includes alignment to the GRCh38 reference (1000 Genomes version) with BWA-MEM (v0.7.15)^86^. For plasma cfDNA, we used a modified alignment pipeline to accommodate adapter trimming after observing increased adapter contaminated reads in cfDNA samples as compared to tumor samples, due to the fact that cfDNA has shorter fragment size, which can lead to R1 and R2 overhang. We therefore used Skewer^87^ for adapter trimming (default settings) and subsequently aligned samples using BWA-MEM (default settings) to the GRCh38 reference (1000 Genomes version). For all samples, duplicate marking and sorting was done using NovoSort MarkDuplicates (v3.08.02), a multi-threaded bam sort/merge tool by Novocraft Technologies; http://www.novocraft.com), followed by indel realignment (done jointly for the tumor and matched germline) and base quality score recalibration using GATK (v4.1.8; https://software.broadinstitute.org/gatk), resulting in a final coordinate sorted bam file per sample. Alignment quality metrics were computed using Picard (v2.23.6; QualityScoreDistribution, MeanQualityByCycle, CollectBaseDistributionByCycle, CollectAlignmentSummaryMetrics, CollectInsertSizeMetrics, CollectGcBiasMetrics) and GATK (average coverage, percentage of mapped and duplicate reads). To specifically assess for sample contamination, we applied Conpair^88^, which validated genetic concordance among the matched germline, tumor and plasma samples, as well as evaluated any inter-individual contamination in the samples. Samples that showed low concordance (<0.99) were excluded from further analysis. Specifically, three preoperative plasma samples from LUAD patients 37, 38 and 39 (described previously in MRDetect^28^) and one set of serially monitored cutaneous melanoma samples from the melanoma patient MSK-55 were rejected from analysis due to low concordance score. We also used read depth skews in copy number neutral plasma regions where available as an additional quality metric (see **Plasma read depth denoising**). Here, we computed sample level Z scores in CNV neutral regions (**Supplementary Table 1**) using our read depth classifier and samples with a Z score value > 10 were excluded. One adenoma plasma sample, Aar-35, was excluded under these criteria. An additional tumor sample, Aar-15, was excluded due to low tumor purity (<30% as assessed by Sequenza^89^, **Supplementary Table 1**), which precluded accurate SNV identification (number of somatic mutations < 1,000, **Supplementary Table 1**) in FFPE tumor tissue (see Tumor / Normal somatic mutation calling).

### Tumor / Normal somatic mutation calling

The primary tumor and matched germline bam files were processed through the NYGC somatic variant calling pipeline^90^. To achieve stringent somatic variant calling, we enforced high-confidence calls. We further excluded variants that were present at any allelic fraction in the matched normal sample. We note that in the case of our LUAD cohort, where tumor purity was lower (**Supplementary Table 1**), fewer overlapping reads between plasma and tumor mutations were available, and adjacent normal with potential tumor contamination was used rather than PBMC, we used the union of calls among mutation callers to broaden read availability. To further broaden read availability in this cohort, we did not enforce paired-read concordance (Supplementary Table 3). To maintain consistency these standards were also applied to our neoadjuvant (Neo) lung cancer cohort. Small deletions and insertions (indels) were excluded.

CNVs, including deletions, amplifications and copy-neutral LOH, were called using Sequenza (v3.0.0)^89^. We only considered CNVs in autosomal regions (chr1-22) of the genome where the size of the CNV was greater than 1.5Mb. Segments with Depth Ratio of 1 were characterized as neutral while those with Depth Ratio in excess of 1 (Depth Ratio >1.2) were selected as amplifications, and Depth Ratios less than 1 (Depth Ratio < 0.8) were selected as deletions. LOH segments, including copy neutral LOH segments, were selected when Minor Copy-number was assigned 0 by Sequenza.

To filter noise in FFPE tumors^58^, we generated a FFPE tumor blacklist to remove any variant site present in 2 or more tumors in our Aarhus University cohort (*n*=35, **Supplementary Table 1**). Only variants with a VAF greater than 0.2 were selected for analysis to exclude variants with minimal supporting reads in FFPE tissue.

### Tumor-informed plasma cfDNA SNV identification

Detection of patient-specific compendia of SNVs was performed by searching the plasma WGS for all sites from the matched patient-tumor compendium with corresponding mutations in the same genomic site and the same substitution. To efficiently identify variants present in the sequencing data, we used a custom Python script (Python version 3.6.8), which uses the pysam module to efficiently extract alignments harboring variants and extracted any read that both uniquely maps to a variant of interest and was in an aligned portion of the read (no clipping or soft masking at the position of the variant). In all plasma samples we removed a subset of variants through the use of a local recurrent artifact plasma ‘blacklist’ filter generated by aggregating pileup SNVs within our plasma WGS database (*n*=239 WGS plasma samples included in the analysis). Variants with a population allele frequency > 4 or more appearances across patients within our plasma sample database were excluded. We generated a similar blacklist across all plasma sequenced at Aarhus University (*n*=50, **Supplementary Table 1**) to account for local artifact bias^91^ and excluded any variants present in 2 or more plasma samples due to the smaller number of samples in this cohort. To further exclude potential germline variants, we used the gnomAD database (version 3.0) which contains genetic variants from >70,000 whole genomes^92^. We downloaded the gnomAD version 3.0 variant call format (VCF) file that was available in hg38 coordinates from the gnomAD browser. We annotated single base changes that we identified with their population allele frequency and removed any candidate variants if the variant was present in gnomAD with an allele frequency > 1/100. Finally, we excluded variants from simple repeat regions and centromeres from a problematic region blacklist^93^.

### Construction of ctDNA SNV training sets and feature space

All training sets were derived from plasma enriched for ctDNA SNV fragments (true label) from specific tumor types and cfDNA SNV fragments (false label) from healthy controls without known cancer processed in the same location and sequenced under the same settings. **Supplementary Table 2** lists samples used in training for LUAD, CRC, and melanoma. To identify informative features, we first implemented quality filters to filter low-quality noise, germline SNPs, and genomic DNA contamination (see **Supplementary Table 3** for quality filters by model type). Broadly, filters focused on removing SNV fragments with low base quality (<25 on Phred scale), low depth (<10 supporting reads), and fragment size within 40 bp - 240 bp to reduce genomic DNA contamination. Germline variants were excluded through filtering high VAF variants (VAF <0.2) except in cases where estimated iChorCNA TF was > 0.2. We further enforced that candidate variants were present on overlapping paired reads.

To maximize the accuracy of true (positive) labels, we devised the following strategies to limit noise contamination in our ctDNA (true label) SNV fragment sets. In all true label settings, we used training samples from patients with high burden metastatic disease (TF 9-24% as called by iChorCNA^10^, **Supplementary Table 2**). In samples where we obtained matched tumor tissue, we nominated ctDNA SNVs by intersecting tumor high confidence somatic calls from the NYGC Somatic Pipeline^90^ with SNVs in plasma. When matched tumor tissue was not available, we called mutations directly in the plasma against normal germline sample using Mutect2^94^, leveraging the high TF in these samples to identify consensus somatic mutations (**Supplementary Table 2**). To further filter noise, when possible we used the intersection of ctDNA SNV fragments from two high TF timepoints from the same patient (**Supplementary Table 2**).

Candidate feature evaluation was performed on SNV fragments after applying quality prefiltering (**Supplementary Table 3**) in both true and false labels. Features and corresponding single variant AUC scores are reported in **Supplementary Table 2**. Several strategies were employed to create tissue-specific regional features that could inform the regional likelihood of somatic mutagenesis. Quantitative features were min / max normalized to values between 0 and 1. To evaluate local tumor mutational density, we aggregated WGS SNV mutation calls from the PCAWG database^81^ and counted the aggregate number of SNV mutations across all available tumor samples in a specific primary disease (e.g. melanoma). Local transcription factor and histone CHiP-Seq marks as well as tissue specific bulk RNA expression values were calculated as reads per kilo base per million mapped reads (RPKM) and were drawn from primary tissue alignments in ENCODE^95^. For each feature category (e.g. H3K4me3 ChIP-Seq marks), we assessed all alignments in ENCODE and selected alignments with the highest Pearson correlation between training set true and false label SNVs on Chromosome 1. In certain cases where strong (>0.15) positive and negative correlations were observed, we included alignments for both positive and negative correlations as separate model features. Regional DNase peaks were downloaded as *narrowpeak* files from ENCODE^95,96^ and lifted to GRCh38. Disease-specific ATAC peak calls were downloaded from TCGA^82^. Plasma WGS sequencing error density was calculated by aggregating all SNV pileup variants from non-cancer control plasma sequenced at the New York Genome Center (Control Cohorts A and C, **Supplementary Table 4**). For each of these features, quantitative values were calculated in a sliding interval window around candidate SNV fragments. The length of this window was optimized by comparing the correlation between feature and label between our training set true and false label SNVs on Chromosome 1 alone. Interval lengths are reported in **Supplementary Table 3**. ChromHMM^83^ chromatin annotation tracks were downloaded from ENCODE and lifted to GRCh38. HI-C compartment information was drawn from Hi-C SNIPER^97^ bed files. Replication timing and mean expression values were drawn from prior work^37^ and lifted to GRCh38. Other features, including distance to bound transcription factor^98^ and SNV distance to nearest nucleosomal dyad in lymphocytes^99^, were drawn from prior work and lifted to GRCh38. **Supplementary Table 3** lists features used in each model type.

### SNV deep learning model architecture and model training

To evaluate SNV fragments with our machine learning architecture, candidate SNV fragments were pulled from alignment files using pysam (v0.15.2) and salient features were encoded as input to our deep learning model architecture (**Fig 1d**) with a custom python (v3.6.8) script. There are two main components of our deep learning SNV model architecture: a regional MLP, and a fragment CNN. The MLP takes a tabular feature representation as input and consists of five fully-connected layers with ReLU activation functions of decreasing size. Each layer is preceded by a batch normalization layer and followed by a dropout layer (with the exception of dropout following the final layer).

We represent cfDNA fragments as an 18×240 tensor (**Fig 1d**). Within the rows of the tensor we compare the one-hot encoded reference sequence to the R1 and R2 sequence of a cfDNA fragment containing a variant (either true somatic mutation or sequencing artifact). We also encode the length and position of R1 and R2, and we mark the position of the SNV to be classified as ctDNA or noise. The columns of the matrix mark individual nucleotides along the length of the fragment. The R1 and R2 regions are padded with neutral values (0.2 in each of the 5 possible nucleotides N, A, C, T, G) where the read does not overlap the reference sequence. This tensor serves as input to a CNN which consists of 4 one dimensional convolution layers (convolving over the base pair width dimension), each followed by a max pooling operation. This is then followed by three fully-connected layers (with ReLU activation) and a subsequent dropout layer, and ends with a single sigmoid-activated fully-connected layer (parallel to the MLP). Model architectures are built in Keras (v.2.3.0) with a Tensorflow base (1.14.0). The fragment tensor has potential access to features including fragment length, key genomic features including mutation type, trinucleotide context, and leading or lagging strand, and quality metrics such as PIR and edit distance (how many variants against the reference sequence are present in a fragment). The tensor structure is coded to account for all possible CIGAR outputs, including insertions, deletions, skips, and soft masks, by inserting ‘N’ (base undetermined) values in reads (deletions, soft skips, soft masks) or the reference sequence and as needed in the alternate read (insertions).

Finally, to integrate fragment and regional information, an ensemble classifier with sigmoid activation jointly evaluates the latent space outputs from both the fragment CNN and regional MLP to generate a score between 0 and 1, reflecting the model-based likelihood that a candidate variant containing cfDNA fragment harbors a true somatic mutation (1) vs. a sequencing artifact (0).

We trained our deep learning classifiers (melanoma, CRC, LUAD) using Keras with tensorflow background on fragments from our disease specific training sets (LUAD, CRC, and melanoma, **Supplementary Table 2**) chosen at the sample level. Validation sets were held out from training and drawn from separate patient samples. All performance metrics, including F1, AUC and accuracy within balanced sets, are reported for training sets and validation sets (**Supplementary Table 2**).

### Comparison of MRD-EDGE SNV deep learning classifier performance to other machine learning models

The MRD-EDGE ensemble classifier (**Fig 1d**) was compared to its individual components (fragment CNN and regional MLP) and other machine learning architectures (MLP and random forest model) by randomly subsampling without replacement in ten parts ctDNA and cfDNA SNV fragments from the held-out melanoma validation set (**Supplementary Table 2**) and assessing F1 performance on each subsampling set (**Extended Data 1b**). To assess fragment-level features in the Random Forest and MLP models, salient features were encoded as tabular values, including one-hot categorical encodings for trinucleotide context and mutation type of the candidate SNV as well as numerical representation of fragment-length, position of the variant within the read (PIR), read 1 length, and read 2 length. The MLP for Fragment + Regional Features has the same architecture as the Regional MLP (see **SNV deep learning model architecture and model training**). The Random Forest Fragment + Regional Features model was constructed using the Python (version 3.6.8) module sklearn sklearn.ensemble.RandomForestClassifier with default settings.

### Generation of synthetic-plasma DNA admixtures

For MRD-EDGE SNV performance evaluations, we generated *in silico* admixtures (range, 10^-7^–10^-3^) from MEL-01 plasma and plasma from a healthy control patient without known cancer (patient C-16). For MRD-EDGE CNV performance evaluations, given the challenges of applying LOH-based classification on samples with different germline SNPs, we generated *in silico* dilutions, with varying fractions (range 10^-6^–10^-3^), of reads from a pretreatment high burden melanoma plasma sample (AD-12 pretreatment timepoint, TF 17% with 1.6 GB of total aneuploidy) into a posttreatment plasma sample from the same patient following a major response to immunotherapy (AD-12 Week 6 Timepoint, TF <5% without observable aneuploidy). We similarly admixed a pre- and postoperative plasma sample from a patient with NSCLC (Neo-03, TF 3.6% with aneuploidy matching tumor CNVs preoperatively, no aneuploidy postoperatively, **Supplementary Table 2**). SAMtools (v1.1, view -s and merge commands) was used to downsample and admix high burden cancer plasma cfDNA reads into low burden (for CNV performance evaluation) or healthy control (for SNV performance evaluation) plasma cfDNA reads accounting for TF and tumor ploidy.

The downsampling ratio S to generate dilutions at various TFs was described previously^28^ and is as follows:

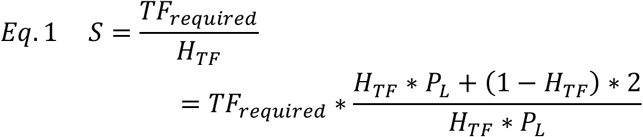

Where H_TF_ denotes ctDNA TF in the high burden cfDNA sample, P_L_ denotes ploidy in the tumor sample. High burden and control coverage is scaled followed by merging of reads:

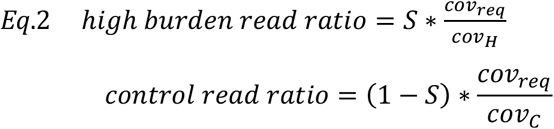

Where cov_req_ is the required read depth coverage for the admixture sample and cov_H_, cov_C_ are the read depth coverage of the high burden and control samples, respectively.

### Plasma SNV-based ctDNA detection and quantification in the tumor-informed approach

As described previously^28^, we modeled the relationship between coverage, mutation load (SNV/tumor), number of detected variants in cfDNA WGS, and the tumor fraction according to the following equation:

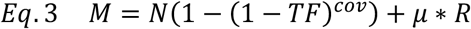

Where *M* denotes the number of SNVs detected in the plasma sample, *N* denotes the number of SNVs (mutation load) in the patient-specific mutational compendium, *TF* denotes the tumor fraction, *cov* denotes the local coverage in sites with a tumor-specific SNV, *μ* denotes the mean noise rate (number of_errors/number of reads evaluated) that corresponds to the patient-specific SNV compendium evaluated in control plasma WGS data (see below), and *R* denotes the total number of reads covering the patient-specific mutational compendium. This relationship allows the calculation of the plasma TF from the mutation detection rate, even in extremely low allele fraction where the mutation allele fraction itself is not informative (random sampling between 0 and 1 supporting read at best).

To address variation in sequencing artifact noise (*μ*) across patients with different mutational compendia, we apply the patient-specific mutational compendium to calculate the expected noise distribution across the cohort of control plasma samples. The process described above is performed to detect the patient-specific SNVs in control plasma samples or other patients (cross-patient analysis). These detections represent the background noise model for which we calculate the mean and standard-deviation (μ,σ) of artifactual mutation detection rate. Confident ctDNA detection can then be defined by converting the patient-specific detection rate (*det_rate* = number of SNVs detected in cfDNA/number of reads checked = M/R) to a 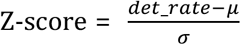, and define a threshold that will keep the specificity above 90%. Specificity and sensitivity performance values were further validated using receiver operating characteristic (ROC) curve using the Python (version 3.6.8) module sklearn sklearn.metrics.roc_curve.

Calculating the patient TF from point mutation detection was then carried out by the following equation (which is an inversion of Eq.3) as described previously^28^:

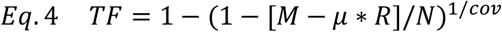

Where *M* denotes the number of SNVs detected in the plasma sample, *N* denotes the number of SNVs (mutation load) in the patient-specific mutational compendium, *TF* denotes the tumor fraction, *cov* denotes the local coverage in sites with a tumor-specific SNV, *μ* denotes the noise rate (number of errors/number of reads evaluated) that corresponds to the patient-specific SNV compendium, and R denotes the total number of reads covering the patient-specific mutational compendium.

### Selection of control plasma samples for tumor-informed approaches

In the tumor-informed setting, patient-specific mutational compendia are applied to both matched plasma and control plasma. To exclude batch specific biases, we employed control plasma samples obtained from the same collection site, sequencing platform and sequencing location as our cancer plasma samples. For example, our early-stage CRC plasma, sequenced at the New York Genome Center on Illumina HiSeq X, was compared to similarly sequenced healthy control plasma (Control Cohort A), while our adenomas and pT1 lesions, sequenced with Illumina NovaSeq 1.5 at Aarhus University in Denmark, was compared to healthy control plasma sourced and sequenced from that institution (Control Cohort B). Control plasma samples used in model training or to construct a read depth classifier PON were not used in downstream analyses (e.g., ROC analyses).

### Plasma read depth denoising

We recently introduced a read depth denoising approach for reducing recurrent noise and bias for WGS-based tumor CNV detection^40^. Our read depth pipeline separates foreground (CNV signal) from background (technical and biological bias) in read depth data by learning a low rank subspace across a panel of normal samples (PON) using robust Principal Component Analysis (rPCA) and applies this subspace to a tumor sample to infer CNV events. To optimize our approach for plasma, we first created PONs from healthy controls plasma generated with the same sequencing preparation (see **Selection of control plasma for tumor-informed approaches, Supplementary Table 3**). We then created log transformed, zero centered read depths across the PON for each sample within 1Kb genomic windows. We performed a window-based rPCA decomposition on our PON to yield a subspace of biases that define “background” noise. Cancer plasma samples were subsequently projected on this background subspace to produce two vectors: a background bias projection and a residual corresponding to plasma CNV read depth skews. We further filtered genomic windows in plasma where read depth was ‘NA’ or was outside of 2.5 standard deviations away from the sample mean.

To generate sample read depth scores for our read depth classifier, we median-normalized window-level read depth values either to sample or chromosome based on mean plasma cohort autocorrelation (to sample < 0.06 < to chromosome, **Supplementary Table 1**). We then aggregated this signal based on the direction of the CNV change in tumor (−1 * deletion and +1 * amplification) to produce a mean per-window read depth score as described previously^28^. This sample level read depth score was compared to read depth scores from held-out control plasma samples in matched genomic regions to generate a final sample-level Z score.

### Plasma CNV-based TF estimation for use in read depth skews

Estimated TFs for our read depth classifier and MRDetect-CNV at different TF admixtures were calculated as:

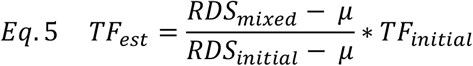

Where RDS_mixed_ is the aggregated median-normalized read depth signal for a specific mixing replicate, RDS_initial_ is the aggregated median-normalized read depth signal for the initial high burden sample, *μ* (noise rate) is the average of aggregated median-normalized read depth signal across held-out plasma controls, and TF_initial_ is the tumor fraction of the initial high burden sample.

### Evaluation of B-allele frequency in plasma

We applied GATK (v3.5.0, https://software.broadinstitute.org/gatk) HaplotyeCaller to identify genome-wide germline SNPs in PBMC WGS data. We then identified major alleles in matched tumor tissue by selecting SNPs with BAF > 0.6 in tumor regions with LOH (see **Tumor / Normal somatic mutation calling**). To enrich for local signal, we grouped SNPs into non-overlapping 1Mb genomic windows. To ensure we evaluated only true SNPs and that our signal was not biased by coverage or subtle clonal mosaicism in PBMCs, we implemented stringent quality filters, including minimal coverage thresholds (plasma and PBMC read depth ≥ 20 reads) and outlier criteria (0.3 < plasma BAF < 0.7, 0.4 < PBMC BAF < 0.6) at the individual SNP level. At the 1Mb window level, we further filtered bins with few SNPs (≤ 50 SNPs/bin) and outlier bins in which the mean plasma or PBMC BAF was outside of 2.5 standard deviations from mean window-level plasma and PBMC BAF from samples sequenced within the same sequencing platform (HiSeq X or NovaSeq). Because 1Mb window-level mean BAF variance is a function of number of SNPs (higher BAF variance with fewer SNPs), we converted window-level BAF values to Z scores normalized for number of window-level SNPs in intervals of 50 SNPs for both plasma and PBMC BAFs, using the range of BAF values for all windows seen in that sequencing platform (HiSeq X or NovaSeq).

Short-read genome sequencing of plasma cannot place SNP variants in phase due to read length limits and the distance between successive SNPs^100^. We faced the technical obstacle of comparing phased variants in cancer plasma samples (identified only through LOH in tumor) to unphased variants in control plasma. To remove the underlying contribution of phasing to aggregate BAF signal, we subtracted window-level PBMC BAF values, where deviations from 0.5 may be due to chance or subtle underlying clonal mosaicism, from window-level plasma BAF values to produce a window-level BAF score that reflects the BAF signal from the contribution of ctDNA in cancer plasma in excess of BAF signal from phased variants alone. In control plasma, where variants cannot be phased, we choose the major allele randomly and aggregate individual SNPs to form window-level BAF noise distributions.

At the sample level, window-level BAF scores are aggregated to produce a mean per-window sample-level BAF score. Sample-level BAF scores in cancer plasma are compared to controls in matching genomic regions to produce a final sample-level Z score that reflects the BAF contribution of ctDNA in cancer plasma compared to matched noise.

### Evaluation of tumor-informed fragment size entropy

We calculated fragment length entropy to capture the heterogeneity of fragment insert size for cfDNA fragments within consecutive non-overlapping 100kb genomic windows. We restricted analyses to fragments with insert size between 100-240bp. First, we calculated in each window the fraction of fragment sizes in each 5bp interval from 100 - 240bp. We then calculated Shannon’s entropy on the set of these fractional inputs. At the sample level, we converted window entropy values from all 100kb windows (neutral and CNV) to median-normalized robust Z scores. By normalizing to the distribution of entropy values in each sample, neutral regions serve as an internal control that accounts for the baseline fragment length heterogeneity within each sample inclusive of entropy noise from different sample preparations and pre-analytic biases. Following normalization, we multiplied window-level Z scores based on the direction of the CNV change using our underlying knowledge of tumor events. We expect more fragment entropy from the contribution of additional ctDNA fragments in tumor amplifications and thus multiply these values by +1, versus less fragment entropy from the contribution of fewer ctDNA fragments in tumor deletions and therefore multiply these values by −1. Regions surrounding transcription start sites (TSS) are known to harbor altered fragmentation profiles including an increase in short fragments^14,44,101^, and this is particularly impactful for regions with deletions in matched tumors, where the shorter TSS fragment signal would confound the anticipated signal of less entropy due to lower contribution of short ctDNA fragments. We therefore excluded bins containing and flanking TSS sites identified in tissue specific ChromHMM^83^ annotations (eg. primary colon TSS for CRC samples) in deletions. We further excluded outlier regions where window-level Z score was greater than 5 median absolute deviations (MADs) from the sample median. We note that recurrent amplifications in chromosome 1p and 22q were uniformly present in control plasma samples in Control Cohort A (*n*=34 plasma samples) and Control Cohort C (*n*=30 plasma samples), and these regions were excluded from analysis as likely cfDNA WGS-specific artifacts.

At the sample level, we aggregated signed window-level CNV Z scores (after multiplication by expected direction based on matched tumor amplification / deletions) across windows to generate a sample-level fragment entropy score. Sample level fragment entropy scores in cancer plasma are compared to controls in matching genomic regions to produce a final sample-level Z score that reflects the contribution of ctDNA in cancer plasma compared to noise in non-cancerous control plasma.

### Removing artifactual CNV events

To reduce CNV artifacts we filtered out genomic bins overlapping centromere and telomere regions (as defined in https://genome.ucsc.edu/ for GRCh38) +/- 5 Mb around each region. Somatic CNV events originating from possible clonal hematopoiesis can also create biases in plasma cfDNA CNV analysis, as most cfDNA is derived from blood cells. To identify such events we evaluated the genome-wide distribution of BAF in PBMC samples as assessed by ascatNgs (v4.2.1) and excluded any regions (variable segment sizes) where the mean BAF was above 0.6. Three patients had detectable somatic PBMC events as described previously^28^: LUAD10 (amp Chr12:60138-133841502), LUAD26 (CN-LOH Chr4:50400000-191044164) and CRC03 (del Chr3:234305- 80851349; del Chr5:75605307-180877637; del Chr7:95649215-125071428; del Chr7:144889607-159128563; del Chr10:50003039-108417985; del Chr15:36365636-63901029; del Chr17:7602691-13317308; del Chr17:17598183 - 20374289; del Chr18:24227106-78017148).

### Aggregation of CNV scores

Our 3 CNV features (read depth, fragment entropy, and BAF) independently inform our estimation of ctDNA signal. We therefore aggregated our features by combining Z scores using Stouffer’s method 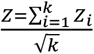.

The MRD-EDGE CNV platform was not applied to our early-stage LUAD cohort due to low tumor purity (median 0.23, range 0.05 - 0.53, 12 / 39 samples with tumor purity ≤ 15%, Supplementary Table 1) which prevented Sequenza from assigning tumor ploidy and total and minor copy number calls in over 30% of samples. Further, in our LUAD cohort, we used adjacent normal tissue rather than PBMC, and therefore we could not assess the underlying PBMC tissue for clonal hematopoiesis events that could serve as a major confounder to our BAF analyses. To assess our neoadjuvant (‘Neo’) NSCLC cohort, we used the same standards as were applied to our LUAD cohort to demonstrate generalizability of our SNV-only approach across sequencing platforms (Illumina HiSeq X in LUAD cohort and Illumina NovaSeq v1.0 in Neo cohort).

For our cohort of adenomas and pT1 lesions, we used our MRD-EDGE SNV classifier to first estimate the TF of detected samples. We found that the estimated TFs of detected lesions by SNV was median 2.88*10^-6^ (range 1.02*10^-6^-1.45*10^-5^) in pT1 lesions and 3.78*10^-6^ (range 1.17*10^-6^-1.21*10^-5^) in adenomas (**Fig 4c**). We therefore reasoned that the LLOD demonstrated in benchmarking for our BAF and fragment entropy CNV features (5*10^-5^) would preclude use in these extremely low TF lesions (**Fig 2c-d**), and indeed our BAF classifier and fragment entropy classifier in these cohorts failed to detect signal in these lesions (AUC 0.51 and 0.48, respectively). We therefore proceeded solely with use of our read depth classifier, which demonstrated sensitivity down to 5*10^-6^ in *in silico* admixtures (**Fig 2b**).

### Integration of SNV and CNV scores

SNV and CNV classifiers provide orthogonal sources of information and are used to independently quantify ctDNA. We evaluated MRD and pT1 / adenoma detection as a sample level Z score in excess of either the CNV or SNV Z score threshold as obtained through calculating the 90% specificity boundary compared to plasma from healthy controls in preoperative early-stage cancer samples. For example, in CRC, we defined a positive detection as a Z score threshold in excess of 90% specificity against healthy control plasma in our preoperative early-stage CRC cohort. We applied these same, prespecified Z score thresholds to identify postoperative MRD (**Fig 3c**) and our pT1 and adenoma lesions (**Fig 4a**). The same was done in lung cancer for our early-stage LUAD and neoadjuvant therapy (‘Neo’) cohorts (**Fig 3d**, **Extended Data 4c**).

### Evaluating SNVs for *de novo* mutation calling

We collected all variants against the hg38 reference genome through samtools (v.3.1) mpileup with no exclusion filters. Only SNVs mapping to chromosomes 1 - 22 were included in our analysis. Indels were excluded. We ran a custom python (v3.6.8) script to collect all fragments containing SNVs that matched pileup variants from the bam alignment. Fragments were then subjected to quality filters and the recurrent artifact blacklist and encoded as inputs to our model architecture (see **SNV deep learning model architecture and model training).** We defined SNV detection rate, a function of the two unknown variables plasma TF and tumor mutational burden (TMB), as the number of fragments classified as ctDNA over the number of post-filter fragments evaluated.

### Determination of *de novo* mutation calling specificity threshold

In a tumor agnostic setting (*de novo* mutation calling), our datasets are more heavily imbalanced between signal and noise than in the tumor-informed setting, where knowledge of tumor SNVs is used to inform candidate variants. We determined the specificity threshold for *de novo* mutation calling within our MRD-EDGE SNV deep learning classifier by optimizing the trade-off at the fragment level between increasing signal enrichment at higher specificity thresholds (**Extended Data 6a**) vs. decreasing signal availability from overly stringent filtering (**Extended Data 6b**). We therefore evaluated performance of our classifier at high specificity thresholds within *in silico* TF admixtures of MEL-01 and a healthy control plasma sample (C-16, Supplementary Table 2). We evaluated detection sensitivity vs TF=0 in admixtures TF=5*10^-5^ and found AUC to be highest at a specificity threshold of 0.995 (**Extended Data 6b**), with decreasing AUC at 0.9975 and 0.9925. We used this empirically chosen specificity threshold for evaluation of plasma TF in subsequent *de novo* mutation calling analyses. We note that the MEL-01 plasma sample used in threshold determination was excluded from all downstream analysis.

### ichorCNA

ichorCNA^10^ (version 2.0) was used as an orthogonal CNA-based method for cfDNA detection and the estimation of plasma TF in high burden plasma samples. We optimized the input setting for more sensitive detection in low-tumor-burden disease using the modified flags -altFracThreshold 0.001, -normal .99 along with a GRCh38 panel of normal (https://gatk.broadinstitute.org/). All other settings were set to default values.

### Tumor-informed and *de novo* targeted panel

MSK-ACCESS^8^ was used as an orthogonal SNV-based method for evaluation of plasma TF in melanoma samples. MSK-ACCESS was run independently on a subset of pre- and posttreatment plasma samples for 14 patients with cutaneous melanoma with available material allowing concurrent analysis. Application of MSK-ACCESS panel and data analysis was performed by the MSK-ACCESS team. Results for the tumor-informed panel were informed by somatic mutations found in matched tumor samples through MSK-IMPACT^102^ and were reported as average adjusted VAF across evaluated genes.

VAF was adjusted to account for copy number alterations at the locus of interest. Copy number alterations are inferred by applying FACETS^103^ to Whole Exome or Whole Genome tumor tissue used in MSK-IMPACT analysis. The ACCESS team assumes that there are no changes to copy numbers of these segments between the IMPACT and ACCESS samples. Adjusted VAF is calculated as follows

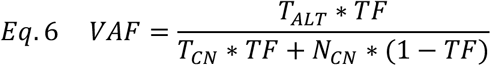

Where *VAF* is the expected variant allele fraction, *TF* is tumor fraction, *T_ALT_* = alternate copies in tumor, *T_CN_* = total copies in tumor, *N_CN_* = total copies in normal.

Solving the equation for *TF* yields:

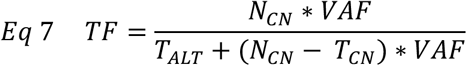

For ACCESS samples, this TF value is computed and named adjusted *VAF* (*VAF_adj_*). For the *de novo* panel, only adjusted VAFs above 0.005 contributed to average VAF.

### Statistical analysis

Statistical analyses were performed with Python 3.6.8 and R version 3.6.1. Continuous variables were compared using Student’s *t*-test, the Wilcoxon rank-sum test or the nonparametric permutation test, as appropriate. All *P* values were two sided and considered significant at the 0.05 level, unless otherwise noted. Cox proportional hazards models were fit using lifelines^104^ and forest plots (**Extended Data 8a**) were plotted using EffectMeasurePlot from zEpid (0.9.0, https://zepid.readthedocs.io/).

## EXTENDED DATA

**Extended Data 1:**
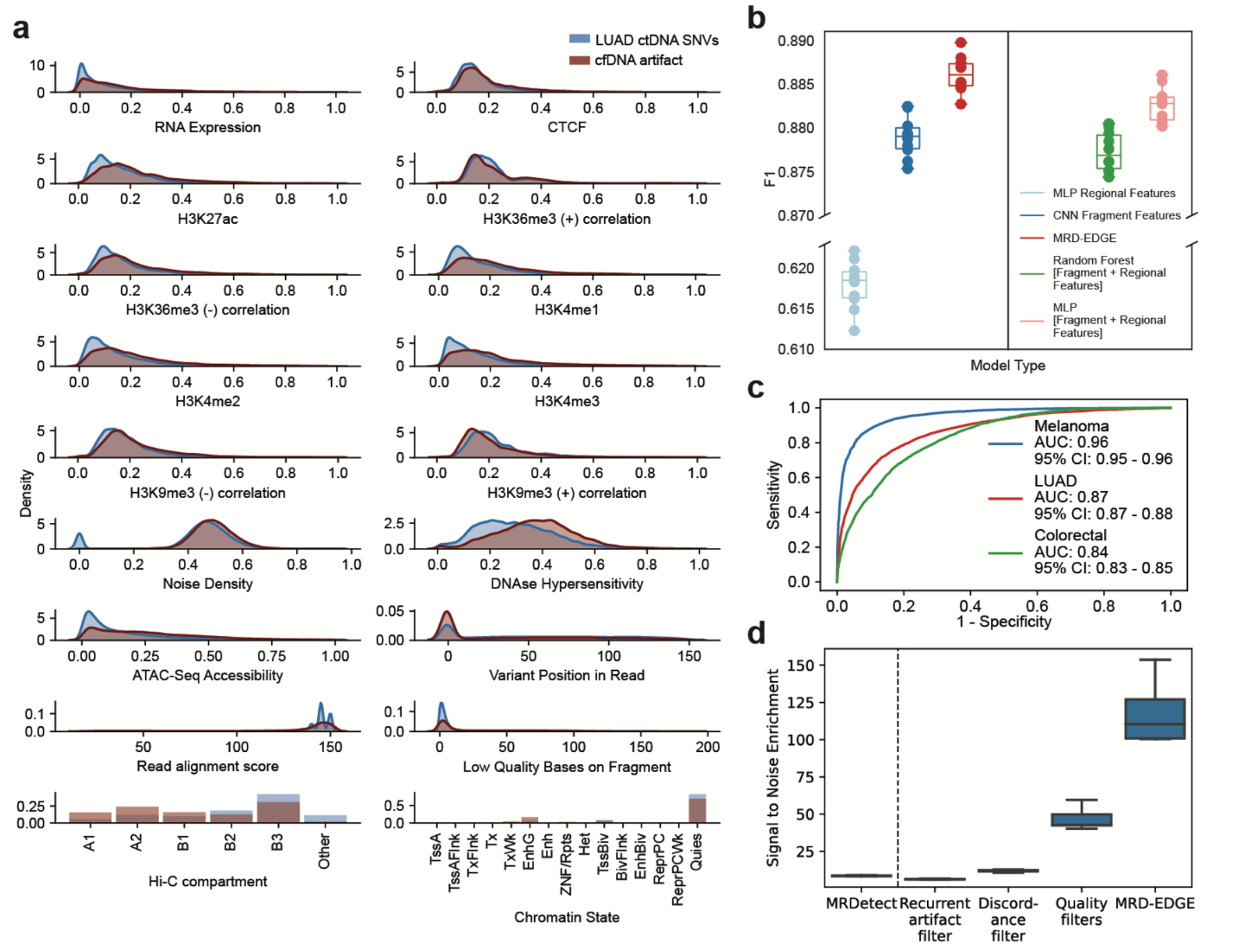
MRD-EDGE SNV feature selection, model architecture and performance. **a)** Feature density plots for post-quality filtered ctDNA and cfDNA SNV artifacts used in the LUAD model. In this comparison, ctDNA SNV fragments are identified from consensus mutation calls in high burden LUAD plasma samples (**Supplementary Table 2**) and cfDNA SNV artifacts are drawn from within the same plasma sample to remove potential inter-sample biases when establishing predictive ability of individual features. **b)** SNV classification performance for different machine learning models. F1 score was assessed on tumor-confirmed melanoma ctDNA SNV fragments vs. cfDNA artifacts from healthy controls. Random subsamplings were drawn from the held-out melanoma validation set (**Supplementary Table 2**), which was split into tenths for this analysis. We compared performance between MRD-EDGE and its separate components (left), as well as to other ML architectures (right) **c)** Fragment-level ROC analysis for MRD-EDGE SNV classifier for different cancer types. Performance is assessed on post-quality filtered fragments (~90% of low-quality cfDNA artifacts are excluded by quality filters) in held-out validation sets (**Supplementary Table 2**) for melanoma, LUAD, and CRC. **d)** Signal to noise enrichment analysis for MRDetect and for each step of the MRD-EDGE tumor-informed pipeline. Final pipeline enrichment is 118-fold for MRD-EDGE vs. 8.3-fold for the MRDetect in the same datasets.

**Extended Data 2:**
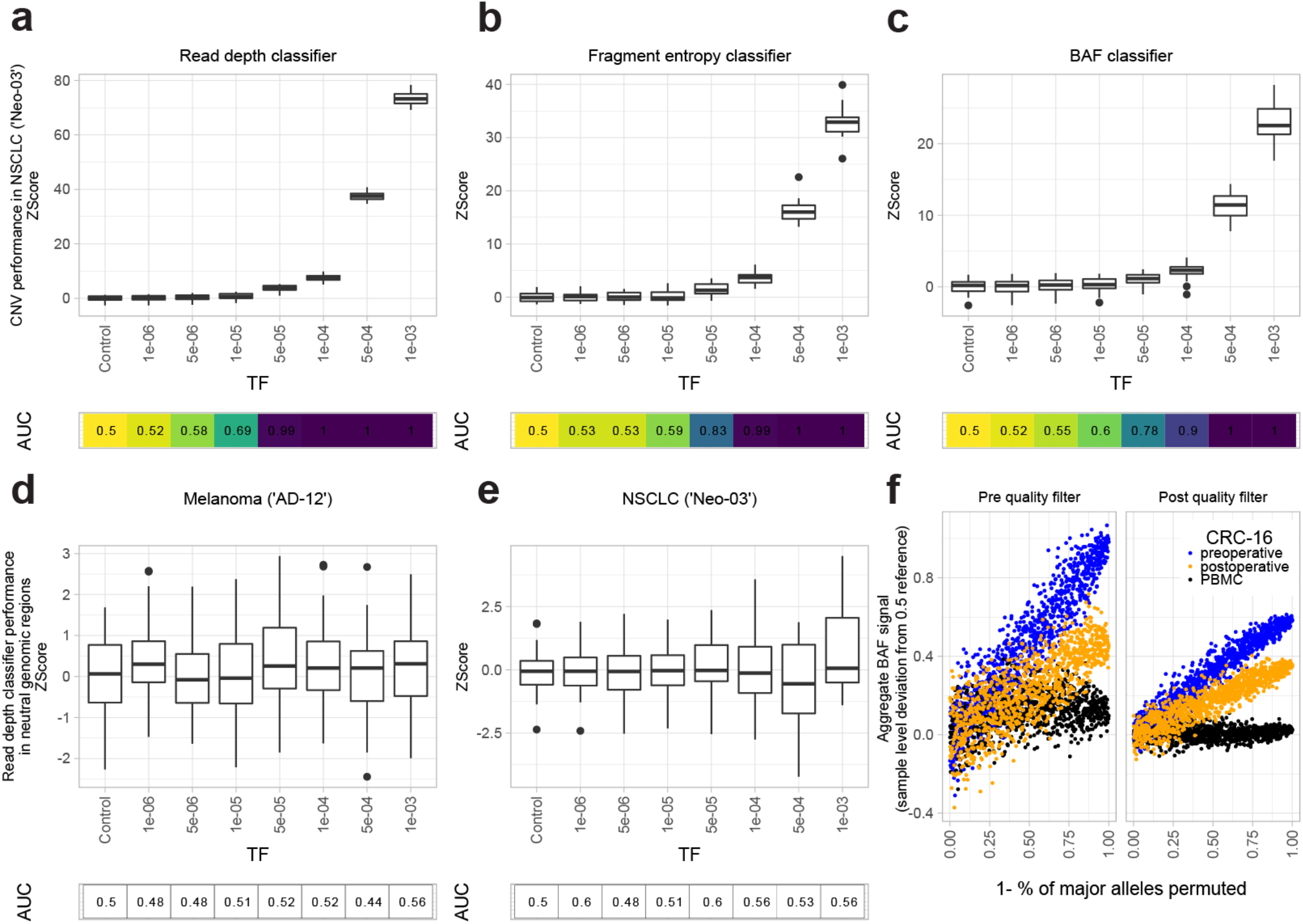
MRD-EDGE CNV detection in neutral regions and non-small cell lung cancer. **a-e**) *In silico* mixing studies in which high TF plasma samples were admixed into low TF samples from the melanoma patient AD-12 and the NSCLC patient Neo-03. For melanoma, pretreatment plasma was mixed into posttreatment plasma as described in **Fig 2b**. For NSCLC, preoperative plasma was mixed into postoperative plasma in 20 technical replicates (each subsampling seed represents a technical replicate). Admixtures model tumor fractions of 10^-6^–10^-3^ (see Methods for detailed description of *in silico* admixture process). Box plots represent median, lower and upper quartiles; whiskers correspond to 1.5 x IQR. An AUC heatmap demonstrates detection performance vs. TF=0 at different mixed TFs as measured by a sample Z score compared to TF=0 distribution for each replicate. The read depth (**a**), fragment entropy (**b**), and SNP BAF (**c**) classifiers demonstrate similar performance in preoperative NSCLC admixtures compared to melanoma admixtures (**Fig 2b-d**). **d-e**, Z scores for the read-depth classifier in neutral regions (no copy number gain or loss in the matched tumor WGS data) for melanoma (**d**) and NSCLC (**e**) demonstrates the expected absence of ctDNA detection at different TF admixtures, consistent with no expected read depth changes in copy neutral regions. **f**) Assessment of preoperative plasma, postoperative plasma, and PBMC BAF in SNPs before (left) and after (right) SNP quality filters in CRC (patient CRC-16). Filters include minimum coverage and outlier exclusion criteria (Methods). BAF signal is calculated as the mean window-level (1Mb) deviation from the 0.5 SNP reference in LOH events identified on matched tumor WGS (Methods), and these values are summed across genome-wide LOH events to calculate sample level signal. To demonstrate the relationship between signal and phased SNPs, the major allele in plasma is randomly permuted to be in phase or out of phase at the percentage specified along the x axis. Following quality filtering, signal can be appropriately inferred and demonstrates the expected relationship between preoperative plasma (highest signal), postoperative MRD (intermediate signal), and PBMC BAF (minimal signal).

**Extended Data 3:**
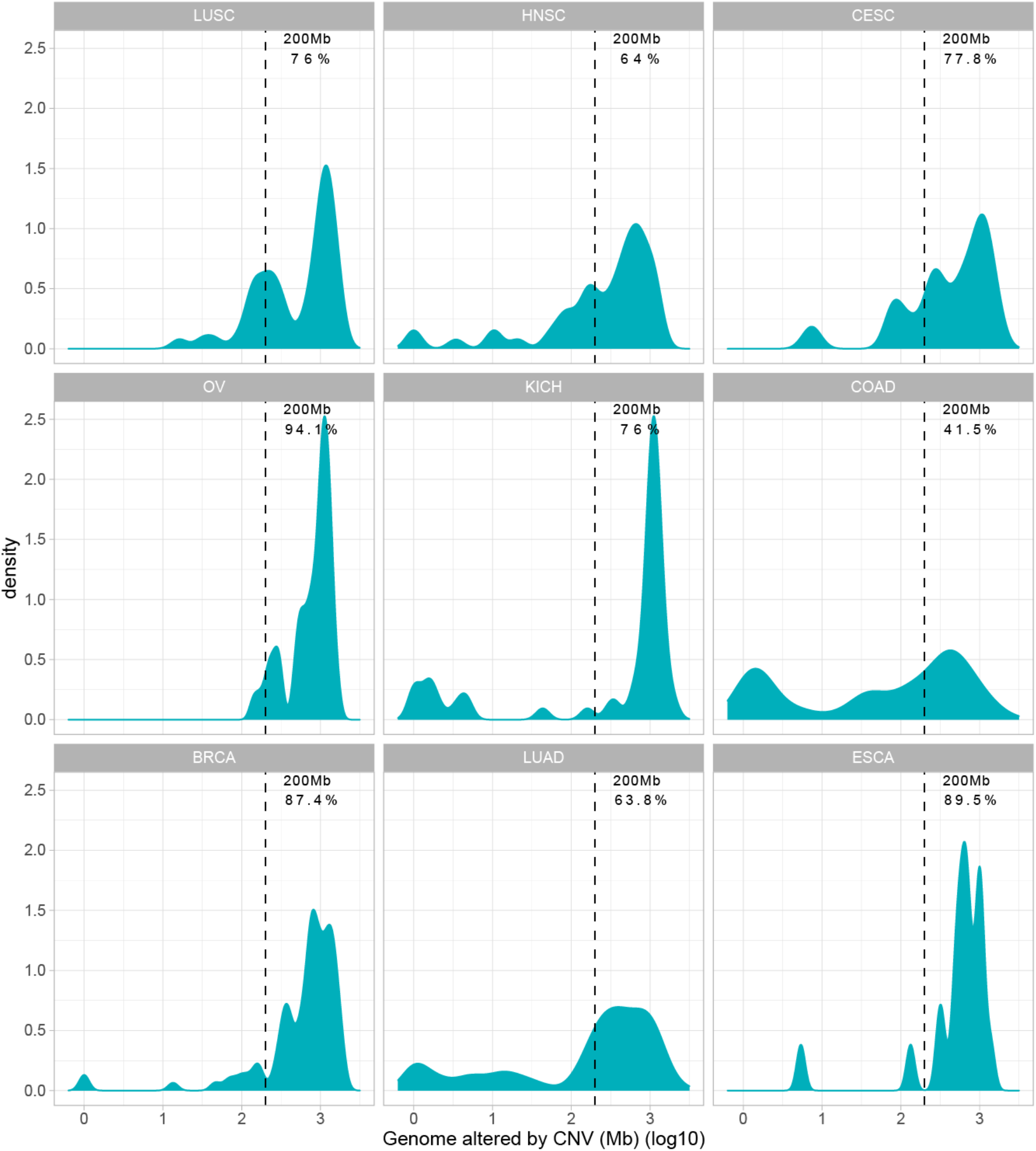
CNV load across tumor types. CNV load in WGS samples across cancer types from the TCGA cohort measured as a function of the size of genome altered by CNV (in loglOMb). Dashed lines represent the percentage of samples that have CNV load of over 200 Mb, the lower limit of detection for the MRD-EDGE CNV classifier. Cancer types include LUSC: Lung squamous cell carcinoma (*n*=50), HNSC: Head and Neck squamous cell carcinoma (*n*=50), CESC: Cervical squamous cell carcinoma and endocervical adenocarcinoma (*n=*18), OV: Ovarian serous cystadenocarcinoma (*n*=50), KICH: Kidney Chromophobe (*n*=50), COAD: Colon adenocarcinoma (n = 53), THCA: Thyroid carcinoma (*n*=50), LUAD: Lung adenocarcinoma (*n*=152), ESCA: Esophageal carcinoma (*n*=19).

**Extended Data 4:**
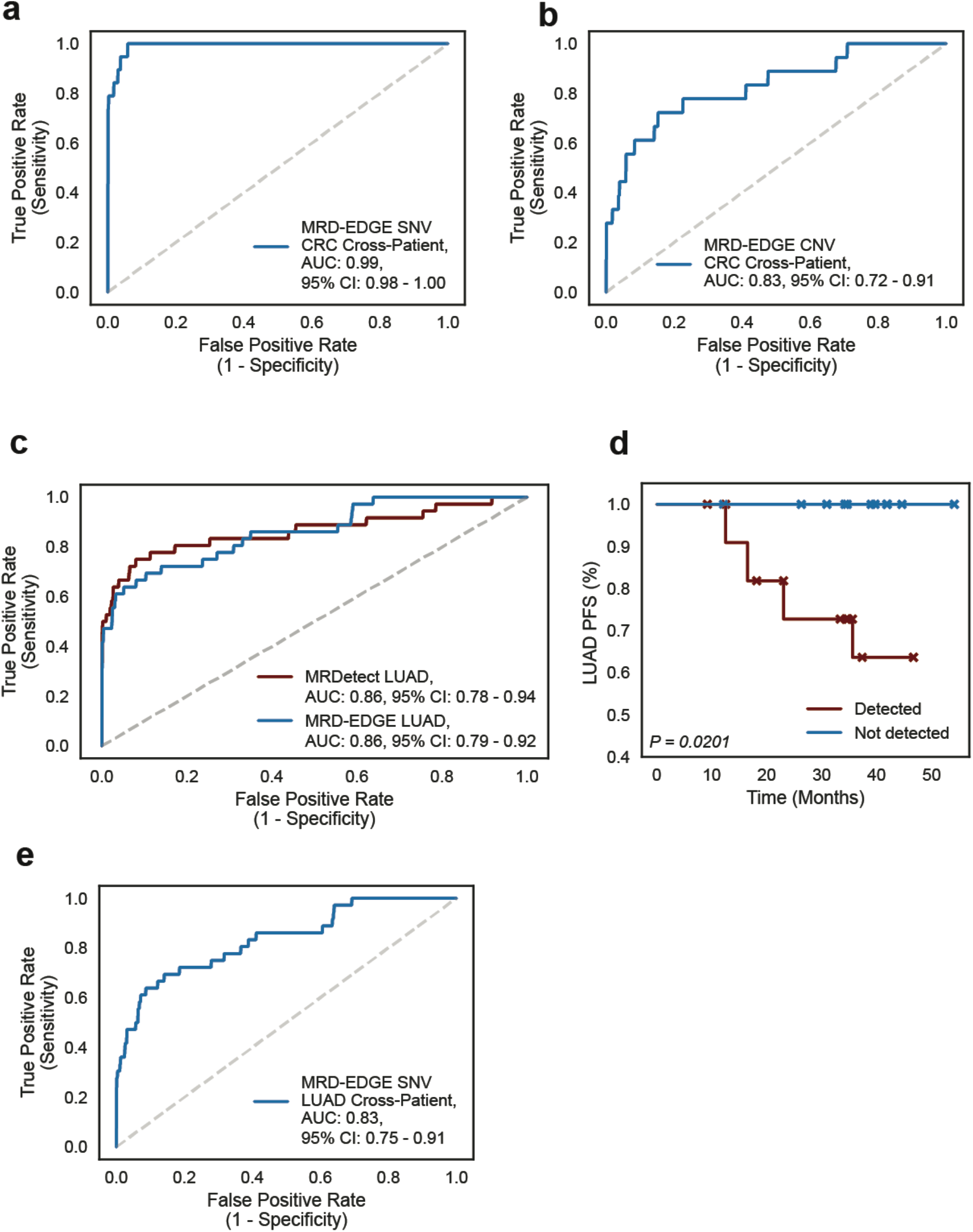
Clinical performance of MRD-EDGE in perioperative CRC and LUAD tumor burden monitoring. **a)** Cross-patient ROC analysis on preoperative colorectal SNV mutational compendia for MRD-EDGE demonstrates similar performance to control (non-cancer) plasma ROC analysis (**Fig 3a**). Preoperative plasma samples (*n*=19) were used as the true label, and SNVs identified from the patient-specific mutational compendia in other preoperative CRC patients (*n*=342; 19 mutational compendia assessed across 18 cross-patient samples) was used as the false label. **b)** Cross-patient ROC analysis on preoperative colorectal CNV mutational compendia for MRD-EDGE. Preoperative plasma samples (*n*=18) were used as the true label, and cross patient plasma was used as the false label (*n*=306; 18 mutational compendia assessed across 17 cross-patient samples) was used as the false label. One sample was excluded due to insufficient aneuploidy. **c)** ROC analysis on preoperative LUAD SNV mutational compendia for MRD-EDGE (blue) and MRDetect SNV + CNV mutational compendia (published previously^28^, red). Preoperative plasma samples (*n*=36) were used as the true label, and the panel of control plasma samples against all patient mutational compendia (*n*=1,224; 36 mutational compendia assessed across 34 control samples from Control Cohort A) was used as the false label. **d)** Kaplan-Meier disease-free survival analysis was done over all LUAD patients with detected (*n*=12) and non-detected (*n*=10) postoperative ctDNA. Postoperative ctDNA detection shows association with shorter recurrence-free survival (two-sided log-rank test). **e)** Cross-patient ROC analysis on LUAD colorectal SNV mutational compendia for MRD-EDGE demonstrates similar performance to control (non-cancer) plasma ROC analysis. Preoperative plasma samples (*n*=36) were used as the true label, and SNVs identified from the patient-specific mutational compendia in other preoperative LUAD patients (*n*=1,260; 36 mutational compendia assessed across 35 cross-patient samples) were used as the false label.

**Extended Data 5:**
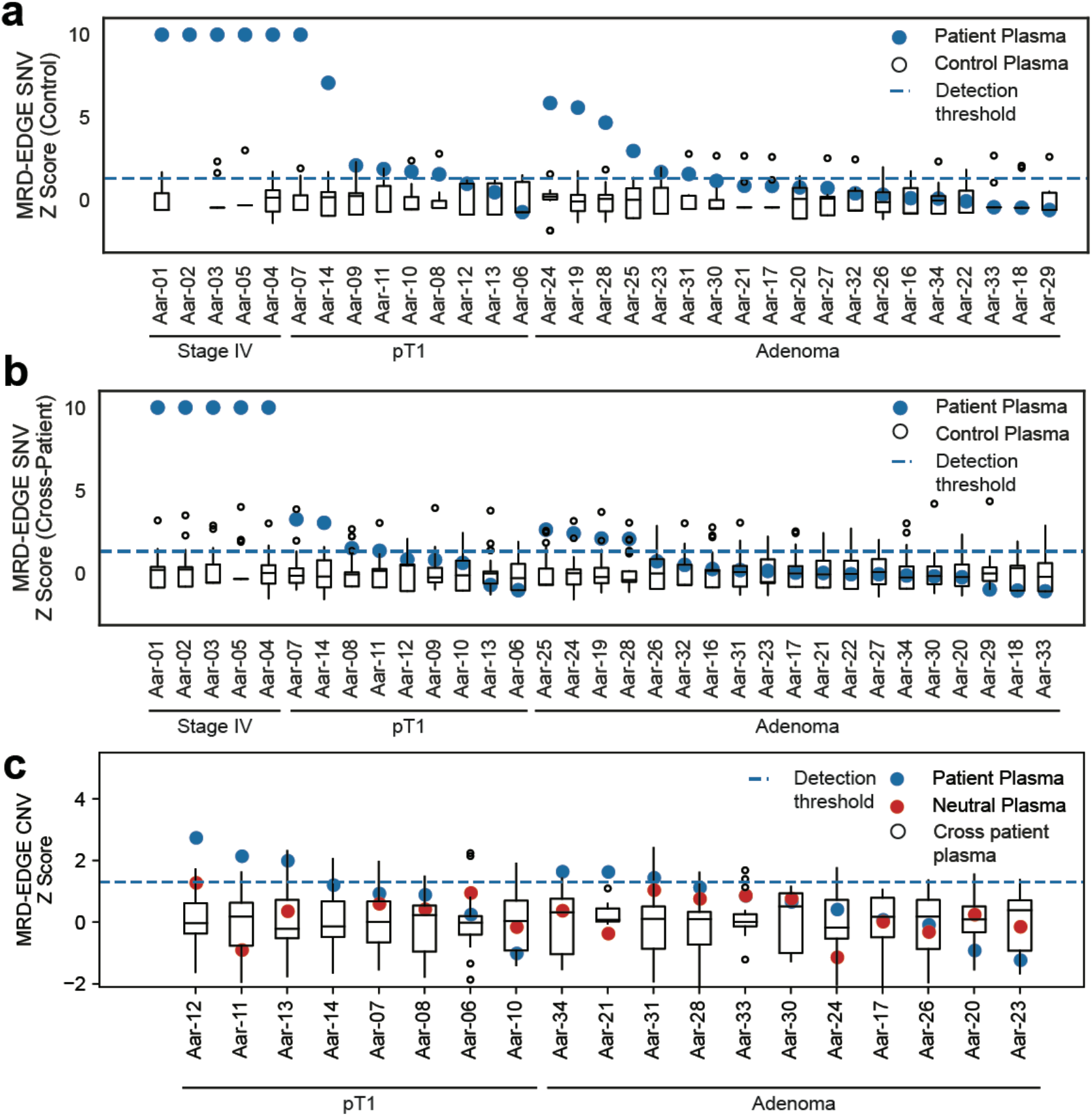
MRD-EDGE detection of ctDNA from colorectal pT1 carcinomas and adenomas. **a)** MRD-EDGE SNV Z score discrimination between signal detected in patient plasma (blue dots, *n* = 33 patients) and healthy control plasma from Control Cohort B (white boxes, *n*=11). Four additional samples from Control Cohort B were used in model training and were therefore excluded from downstream SNV analysis. Signal is measured on patient plasma and the control plasma samples using the same patient-specific SNV compendium. The SNV ctDNA detection threshold (dashed horizontal line) was prespecified, reflecting 90% specificity defined in an independent cohort of preoperative patients with early-stage CRC (**Fig 3a**). **b**) Cross patient SNV evaluation. SNV Z-score discrimination is calculated as in (**a**) using cross-patient evaluation instead of healthy control plasma. Cross-patient signal is calculated via application of the patient-specific mutational compendium to all other patient plasma (white boxes, *n*=32). The ctDNA detection threshold (dashed horizontal line) was prespecified, reflecting 90% specificity defined in an independent cohort of preoperative patients with early-stage CRC (**Fig 3a**). **c)** Z-score discrimination between MRD-EDGE CNV on patient plasma (blue, n = 19 patients) compared to signal detected in neutral regions (as a negative control, red), and cross-patient cohort (n = 18, white). Z-score was calculated using the noise parameters estimated by the control plasma cohort. Samples not evaluated due to insufficient aneuploidy (*n*=9) and samples from Stage IV patients (*n*=5) were excluded from analysis, the latter due to a sparsity of neutral regions in these advanced cancer samples. The CNV ctDNA detection threshold (dashed horizontal line) was prespecified, reflecting 90% specificity defined in an independent cohort of preoperative patients with early-stage CRC (**Fig 3b**).

**Extended Data 6:**
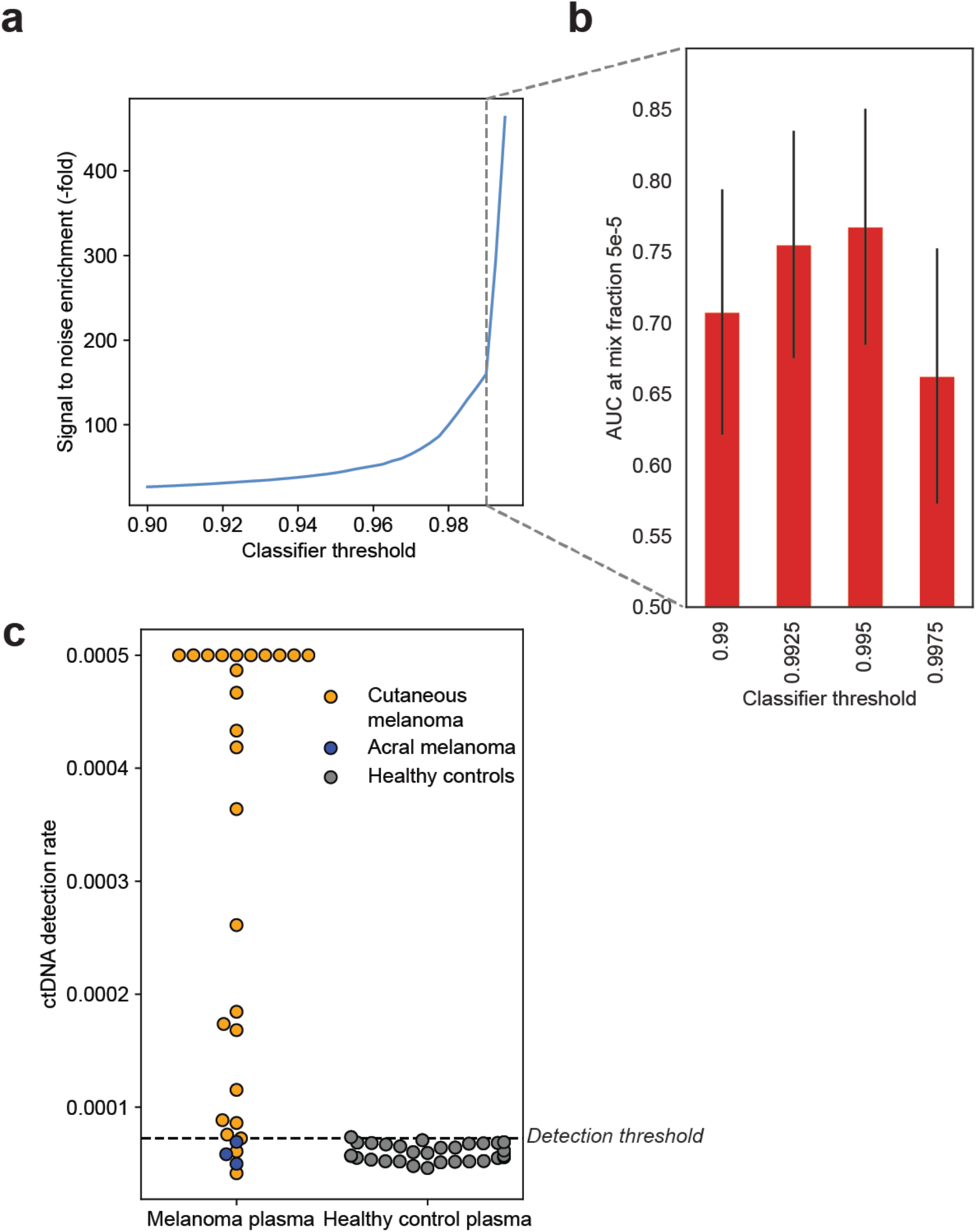
Determination of MRD-EDGE *de novo* mutation calling classification threshold. **a)** Fragment-level signal to noise enrichment, defined as the fraction of remaining ctDNA fragments (signal) over remaining cfDNA SNV artifacts (noise), for different MRD-EDGE classification thresholds in the melanoma held-out validation set derived from tumor-confirmed ctDNA SNVs from the melanoma patient MEL-01 and post-quality filtered cfDNA artifacts from healthy control plasma (**Supplementary Table 2**). The MRD-EDGE SNV deep learning classifier uses a sigmoid activation function that outputs the likelihood between 0 and 1 that a candidate SNV fragment is a mutated ctDNA fragment or cfDNA harboring a sequencing error, and the classification threshold is used as a decision boundary for these two classes. Signal to noise enrichment increases at higher classification thresholds, as expected. **b)** As increased specificity will ultimately eliminate most of the signal, to choose an optimal threshold for classification, we compared sensitivity vs. TF=0 in an *in silico* study of cfDNA from the metastatic melanoma sample MEL-01 mixed in *n*=20 replicates against cfDNA from a healthy plasma sample (TF=0) at 5 * 10^-5^ at 16X coverage depth. We found optimal performance at a classifier threshold of 0.995 as measured by AUC of mixed replicates against TF=0. This threshold was subsequently applied in *de novo* mutation calling analyses. **c)** (left) ctDNA detection rates for pretreatment cutaneous melanoma samples from the adaptive dosing cohort (*n*=26, orange, detection rate was capped at 0.0005) compared to acral melanoma samples (*n*=3, blue, pre- and posttreatment timepoints from 1 patient with acral melanoma) sequenced within the same batch and flow cell. (right) ctDNA detection rates for healthy control plasma (*n*=30, gray). ctDNA is not detected from acral melanoma plasma, demonstrating absence of batch effect and the specificity of MRD-EDGE for the UV signatures associated specifically with cutaneous melanoma.

**Extended Data 7:**
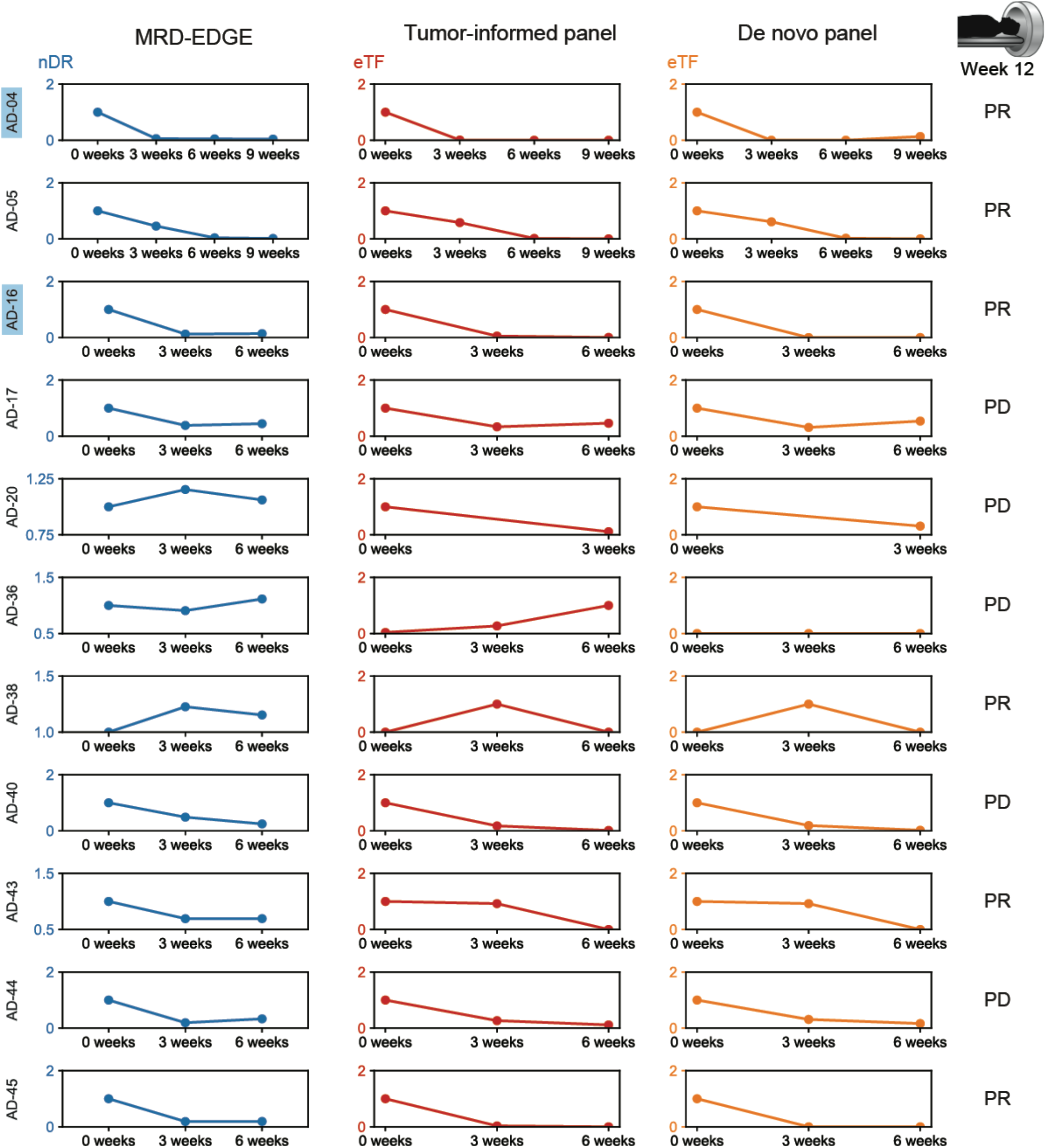
Trends in plasma TF using MRD-EDGE, a tumor-informed panel, and a *de novo* panel. Serial tumor burden monitoring on ICI with MRD-EDGE, tumor-informed panel, and *de novo* panel for 11 patients with melanoma (see **Fig 5f** for remaining 3 patients with matched WGS and panel data). Tumor burden estimates are measured as a detection rate normalized to the pretreatment sample (normalized detection rate, nDR) for MRD-EDGE and as variant allele fraction (VAF) normalized to the pretreatment VAF (normalized VAF, nVAF) in the tumor-informed panel and *de novo* panel. Outcome is reported as RECIST response on Week 12 CT imaging including partial response (‘PR’), stable disease (‘SD’), or progressive disease (‘PD’). Blue highlights surrounding sample names indicate samples with 14 or more mutations covered in the tumor-informed panel.

**Extended Data 8:**
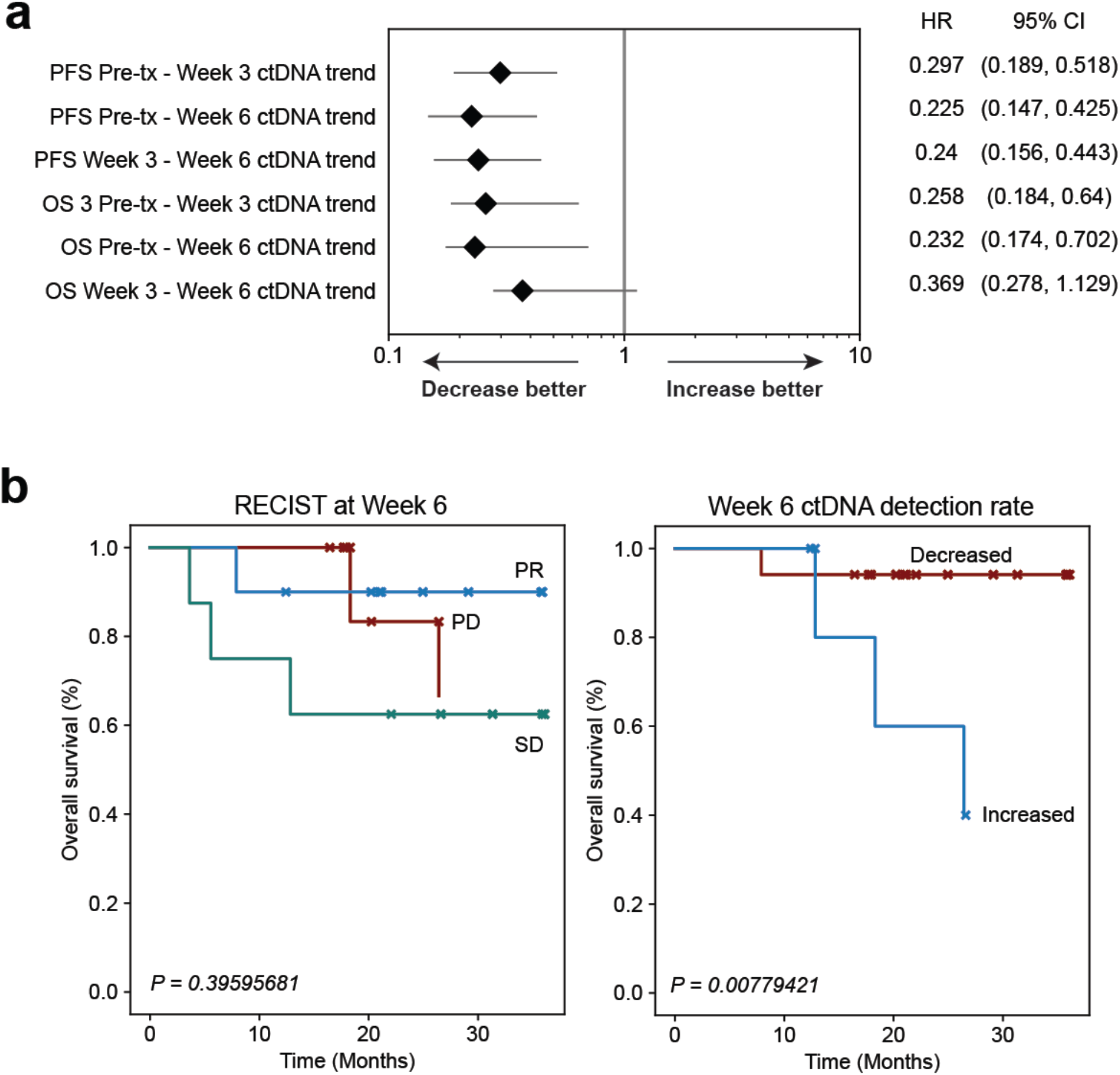
Monitoring response to immunotherapy with MRD-EDGE. **a)** Forest plot demonstrating relationship between ctDNA TF trend (increase or decrease) and progression-free survival (PFS) and overall survival (OS) at serial posttreatment timepoints. MRD-EDGE TF estimates are measured as a detection rate normalized to the pretreatment sample (normalized detection rate, nDR). Each posttreatment timepoint is prognostic of PFS outcomes. **b)** (left) Kaplan-Meier overall survival analysis for Week 6 RECIST response (*n*=10 partial response, ‘PR’, *n=8* stable disease, ‘SD’, *n=6* progressive disease, ‘PD’) in the adaptive dosing melanoma cohort (*n*=26 patients) where CT imaging was available at Week 6 shows no significant relationship with OS (multivariate logrank test). **c)** Kaplan-Meier OS analysis for Week 6 ctDNA trend in adaptive dosing melanoma patients with decreased (*n*=17) or increased (*n*=5) nDR compared to pretreatment timepoint as measured by MRD-EDGE. Patients with undetectable pretreatment ctDNA (*n*=2) were excluded from the analysis as were 2 patients where Week 6 plasma was not available for analysis. Increased nDR at Week 6 shows association with shorter overall survival (two-sided log-rank test). TF, tumor fraction; CT, computed tomography.

